# Pathogenic, but not non-pathogenic, *Rickettsia* evade inflammasome-dependent IL-1 responses to establish an intracytosolic replication niche

**DOI:** 10.1101/2021.09.08.459437

**Authors:** Oliver H. Voss, Jennifer Cobb, Hodalis Gaytan, Natalia Rivera Díaz, Rigoberto Sanchez, Louis DeTolla, M. Sayeedur Rahman, Abdu F. Azad

## Abstract

*Rickettsia* species (spp.) are strict obligate intracellular bacteria, with some being pathogenic in their mammalian host, including humans. One critical feature of these stealthy group of pathogens is their ability to manipulate hostile cytosolic environments to their benefits. Although our understanding of *Rickettsia* cell biology and pathogenesis are evolving, the mechanisms by which pathogenic *Rickettsia* spp. evade host innate immune detection remains elusive. Here, we showed that disease severity in wild-type (*WT*) C57BL/6J mice infected with *R. typhi* (etiologic agent of murine *typhus*) and *R. rickettsii* (etiologic agent of Rocky Mountain Spotted Fever), but not with non-pathogenic *R. montanensis*, correlated with levels of bacterial burden as detected in the spleens, as well as the serum concentrations of pro-inflammatory cytokine IL-1α and to a lesser extent IL- 1β. Antibody-mediated neutralization of IL-1α confirmed a key role in controlling mortality rates and bacterial burdens of rickettsiae-infected *WT* mice. As macrophages are a primary source of both IL-1α and IL-1β cytokines, we determined the mechanism of the anti-rickettsial activities using bone-marrow-derived macrophages. We found that pathogenic *R. typhi* and *R. rickettsii*, but not non-pathogenic *R. montanensis*, eluded pro- IL-1α induction and benefited pre-dominantly from the reduced IL-1α secretion, via a Caspase-11-Gsdmd-dependent pathway, to facilitate intracytosolic replication. Adoptative transfer experiments identified that IL-1α secretion by macrophages was critical for controlling rickettsiosis in *WT* mice. In sum, we identified a previously unappreciated pathway by which pathogenic, unlike non-pathogenic, rickettsiae preferentially target the Caspase-11-Gsdmd-IL-1α signaling axis in macrophages thus supporting their replication within the host.

**IMPORTANCE:** Currently, no vaccines are available to prevent rickettsioses, while vector-borne rickettsial infections in humans are on the rise globally. In fact, the insufficient understanding of how pathogenic *Rickettsia* species circumvent host immune defense mechanisms has significantly hindered the development of more effective therapeutics. Here, we identified a previously unappreciated role for the Caspase-11-Gsdmd-IL-1α signaling axis, to limiting the replication of pathogenic *R. rickettsia* and *R. typhi* species in murine macrophages and wild-type (*WT*) C57BL/6J mice. Adoptative transfer studies further identified IL-1α-secreting macrophages as critical mediators in controlling rickettsial infection in *WT* mice. Collectively, these findings provide insight into the potential mechanism of how pathogenic, but not non-pathogenic *Rickettsia* spp., benefit from a reduction in the Caspase-11-Gsdmd-mediated release of IL-1α to support host colonization.

## INTRODUCTION

Invasive cytosolic bacteria, including *Listeria*, *Shigella*, *Burkholderia*, *Francisella*, *Orientia*, and *Rickettsia* species (spp.) have developed strategies to induce their own uptake by phagocytosis and to circumvent host innate immune defenses for their intracellular survival (1, 2). Human infection with *Rickettsia* spp. occurs via infected hematophagous arthropods such as fleas, ticks, and human body lice (3), either through the bite or deposited infected feces into skin and mucosal surfaces. Upon entry, *Rickettsia* spp. encounter tissue-resident immune cells, like macrophages (MΦ). Activated MΦ play a crucial role in either terminating an infection at an early stage, which commonly is the fate of non-pathogenic *Rickettsia* spp., or succumbing to bacterial replication and pathogen colonization as well as host dissemination to distant organs (3). After internalization into host cells, *Rickettsia* spp. escape from phagosomes and subvert host cytosolic defense mechanisms (*i.e.*, autophagy and inflammasomes) to establish an intracytosolic replication niche. Recently, we reported that pathogenic *Rickettsia* spp. secret effectors to promote host colonization by modulating endoplasmic reticulum structures or by hijacking the autophagic defense pathway (4–9). Subversion of autolysosomal destruction to colonize the host cytosol exposes *Rickettsia* spp. to another cytosolic host sensor-regulated defense mechanism, the inflammasomes (1, 10). Inflammasomes are immune signaling complexes categorized into canonical [Caspase (Casp)-1] and non-canonical [murine Casp-11 or human Casp-4/5] inflammasomes. The inflammasome complex assembly involves the adaptor protein ASC and upstream sensors including NLRP1, NLRP3, NLRC4, AIM2, and pyrin, which are primed by exogenous pathogen-associated molecular pattern molecules (PAMPs) and activated through endogenous damage-associated molecular pattern molecules (DAMPs). Initiation of the canonical inflammasome results in the activation of Casp-1. Active Casp- 1 leads to the maturation of proinflammatory cytokines IL-1β and IL-18 and the activation of gasdermin D (Gsdmd), the executor of pyroptosis (11, 12). Of note, recent findings have also suggested that active Casp-11 is capable of activating Gsdmd (11, 12). Although IL-1β is released by both canonical and non-canonical inflammasome pathways, IL-1α is preferentially released by the non-canonical inflammasome pathway (13–15). IL-1α is expressed by a wide range of hematopoietic and non-hematopoietic cell types, whereas IL-1β is primarily produced by myeloid cells (15). Importantly, although IL-1α and IL-1β signal through the same receptor, IL-1R, these two cytokines are not completely functionally redundant (15). Given the importance for both canonical and non-canonical inflammasome-mediated IL-1 signaling in limiting pathogen colonization, many bacteria have evolved strategies to block their activation (16–21). In fact, various pathogenic intracellular bacteria utilize their own effector repertoire to evade these pathways to successfully colonize and disseminate in their host cells (10, 22).

In the case of *Rickettsia*, our understanding of the role of inflammasomes in controlling host colonization is only now emerging (23–25). Specifically, how pathogenic *Rickettsia* spp. manipulate immune defenses to replicate within the host cytosol not only relies almost exclusively on data from tick-transmitted rickettsiae [*e.g.*, members of spotted fever group (SFG) or transition group (TRG)], but also shows that these pathogens likely employ species-specific strategies to evade host cytosolic defense mechanisms. For instance, *R. australis*, a pathogenic TRG member, benefited from ATG5-mediated autophagy induction and suppression of inflammasome-dependent IL-1β production to colonize both BMDMΦ (23) and mice (26). In contrast, *R. parkeri*, a mildly pathogenic member of SFG (see Fig. 1A), utilizes its surface cell antigen Sca5 (OmpB) for protection against autophagic recognition, and consequently benefits from inflammasome-mediated host cell death that antagonizes the action of Type I IFN in BMDMΦ and mice (24, 27). In contrast, our recent report on the flea-transmitted *R. typhi* [pathogenic member of typhus group (TG)], showed that *R. typhi* is ubiquitinated upon host entry and escapes autolysosomal fusion to establish an intracytosolic niche in non-phagocytic cells (9).

**Fig. 1.**
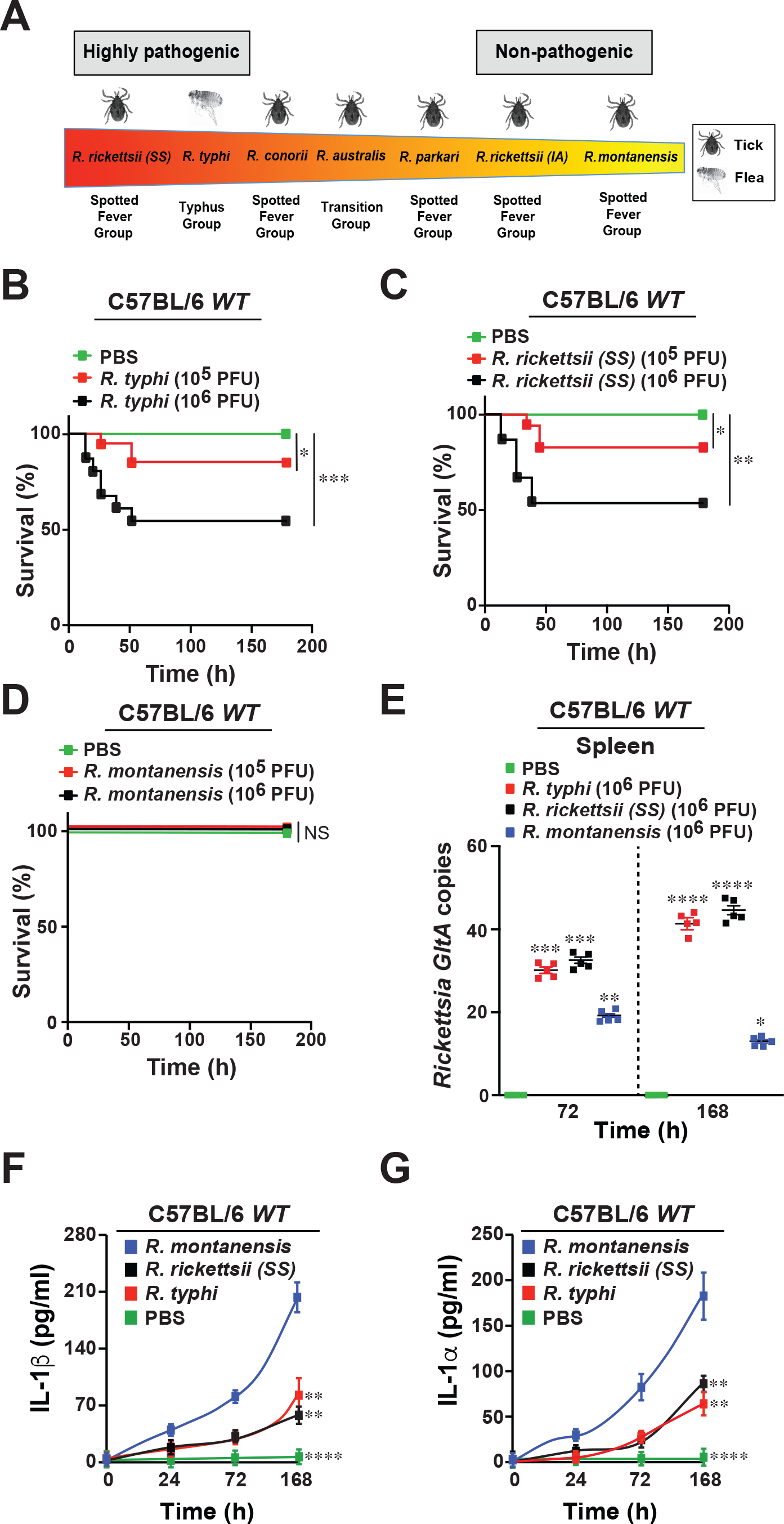
I*n vivo* models of rickettsiosis. (**A**) *Rickettsia* spp. and their level of pathogenicity to humans (3, 43). (**B-D**) Establishment of a model of rickettsiosis (LD25 and LD50) for *R. typhi* [B]*, R. rickettsii* [C] or *R. montanensis* [D] in C57BL/6J *WT* mice. Animals were injected via tail vein (*i.v.*) with different doses [10^5^-10^6^ PFU] of *R. typhi*, *R. rickettsii*, *R. montanensis*, or PBS (n=12 for each treatment). Survival was monitored for 7 days. (**E**) Bacterial burden was tested in spleens of *R. typhi*-, *R. rickettsii*-, *R. montanensis*-, or PBS-injected *WT* mice (n=5 per each treatment) shown in (B-D) by *GltA* RT-qPCR at day 3 and 7 (n=5 for each treatment) using the host housekeeping gene, *Gapdh*, for normalization. (**F-G**) Serum samples from rickettsiae-infected mice described in (B-D) were analyzed for IL-1β (F) and IL-1α (G) production using the Legendplex kits (BioLegend) followed by flow cytometry analysis. Error bars in (E-G) represent means ± SEM from five independent experiments; NS, non- significant; \**p* ≤ 0.05; \*\**p* ≤ 0.01; \*\*\**p* ≤ 0.005; \*\*\*\**p* ≤ 0.001.

These unexpected findings on how members of SFG, TRG, and TG *Rickettsia* differentially promote intracytosolic host survival prompted us to explore the underlying mechanism(s) by which pathogenic, but not non-pathogenic *Rickettsia* spp., block immune defense responses to establish a replication niche in phagocytic cells, like MΦ. Specifically, we sought to test the hypothesis that pathogenic, but not non-pathogenic, *Rickettsia* spp. reduce inflammasome-mediated IL-1 responses thereby promoting their intracytosolic replication within host cells.

## RESULTS

### *In vivo* models of rickettsiosis

At present, more than 30 *Rickettsia* spp. have been described globally, but less than a dozen is known to cause disease in humans, with some being notoriously pathogenic and associated with high morbidity and mortality, while others exert limited or no pathogenicity (Fig. 1A). We sought to test our hypothesis that pathogenic, but not non-pathogenic, *Rickettsia* spp. evade immune responses in host defense cells, like MΦ, to replicate and disseminate. We simultaneously evaluated the cytosolic host defense responses between pathogenic [*R. rickettsii* (*Sheila Smith, SS*) and *R. typhi* (Wilmington) and non-pathogenic (*R. montanensis*) strains *in vivo*. We first established a mouse model of mild (∼LD25) or more severe (∼LD50) rickettsiosis in C57BL/6J *WT* mice. For both *R. rickettsii* and *R. typhi* LD25 or LD50 were achieved with doses of 10^5^ or 10^6^ PFU, respectively (Figs. 1B-C), however, *R. montanensis*-infected mice showed no signs of lethality at either 10^5^ or 10^6^ PFU (Fig. 1D). Bacterial burden in spleens from infected C57BL/6J *WT* mice (only dose of 10^6^ PFU is shown) confirmed successful infection with all three *Rickettsia* spp. at day 3 post-infection, while *R. typhi*- and *R. rickettsii*-infected *WT* mice displayed significantly higher bacterial burden in the spleens as compared to splenic tissues from *R. montanensis*-infected mice at day 7 (Fig. 1E). This correlated with the observed differences in the spleen sizes and weights of the infected animals (Fig. S1). Given the earlier findings from other laboratories and ours (9, 23–28), we hypothesized that the observed dissimilarities in pathogenicity among *Rickettsia* spp. is likely linked to differences in host defense responses. Recent findings further suggest that the intracytosolic survival of different *Rickettsia* spp. is either supported or suppressed by immune defense responses (*e.g.*, IFN-I, TNF-α, or IL-1β) (23–28), and thus leaving the precise mechanism to be determined. Therefore, we first sought to evaluate immune defense responses at the level of IL-1β and IL-1α cytokine secretion in the sera of *R. typhi*-, *R. rickettsii*-, and *R. montanensis*-infected animals. The increase in mortality and elevated bacterial burden correlated with reduced serum levels of both IL- 1β and IL-1α cytokines (Figs. 1F-G), suggesting that reduced activation of both IL-1 signaling responses is a potential mechanism for lethality and survival of pathogenic *Rickettsia* spp.

### Anti-rickettsial activity of IL-1α is involved in restricting *Rickettsia* infection

To characterize further the role of IL-1α and IL-1β cytokines in restricting non-pathogenic and pathogenic *Rickettsia* spp. *in vivo*, IL-1α or IL-1β function was neutralized via tail vein (*i.v.*) injection, with anti(α)-IL-1α, αIL-1β, or αIgG-isotype control antibodies (Abs) into C57BL/6J *WT* mice in our established model of mild rickettsiosis (LD25; ∼10^5^ PFU, Figs. 1B-D). Neutralization of IL-1α, and to a much lesser extent IL-1β, was associated with a significant increase in the mortality of *R. typhi*-, *R. rickettsii*-, and *R. montanensis*-infected mice (Figs. 2A-C) and resulted in the development of splenomegaly (Fig. S2), which correlated with an increase in bacterial loads in the spleen (Fig. 2D). The efficiency of Ab-mediated blocking was confirmed by measuring the levels of IL-1β and IL-1α cytokine in the sera of the rickettsiae-infected mice (Figs. 2E-F). Next, we sought to determine the effect of administering recombinant (r)IL-1α or rIL-1β proteins on *Rickettsia* colonization *in vivo*. Accordingly, we administered (*i.v.*) endotoxin-free rIL-1α and rIL-1β proteins following infection with *R. typhi*, *R. rickettsii*, and *R. montanensis*. Pretreatment of mice with rIL-1α, and to a lesser extent rIL-1β, protected C57BL/6J *WT* mice from pathogenic *Rickettsia*-induced lethality (Figs. 3A-C), with both reduced splenomegaly (Fig. S3), and decreased splenic bacterial burdens (Fig. 3D). Moreover, the observed phenotypes correlated with the measured IL-1α and IL-1β serum concentrations (Figs. 3E-F). Collectively, these findings suggest that IL-1α, and to a lesser extent, IL-1β, are involved in restricting the replication and colonization of non-pathogenic and pathogenic *Rickettsia* spp. in C57BL/6J *WT* mice.

**Fig. 2.**
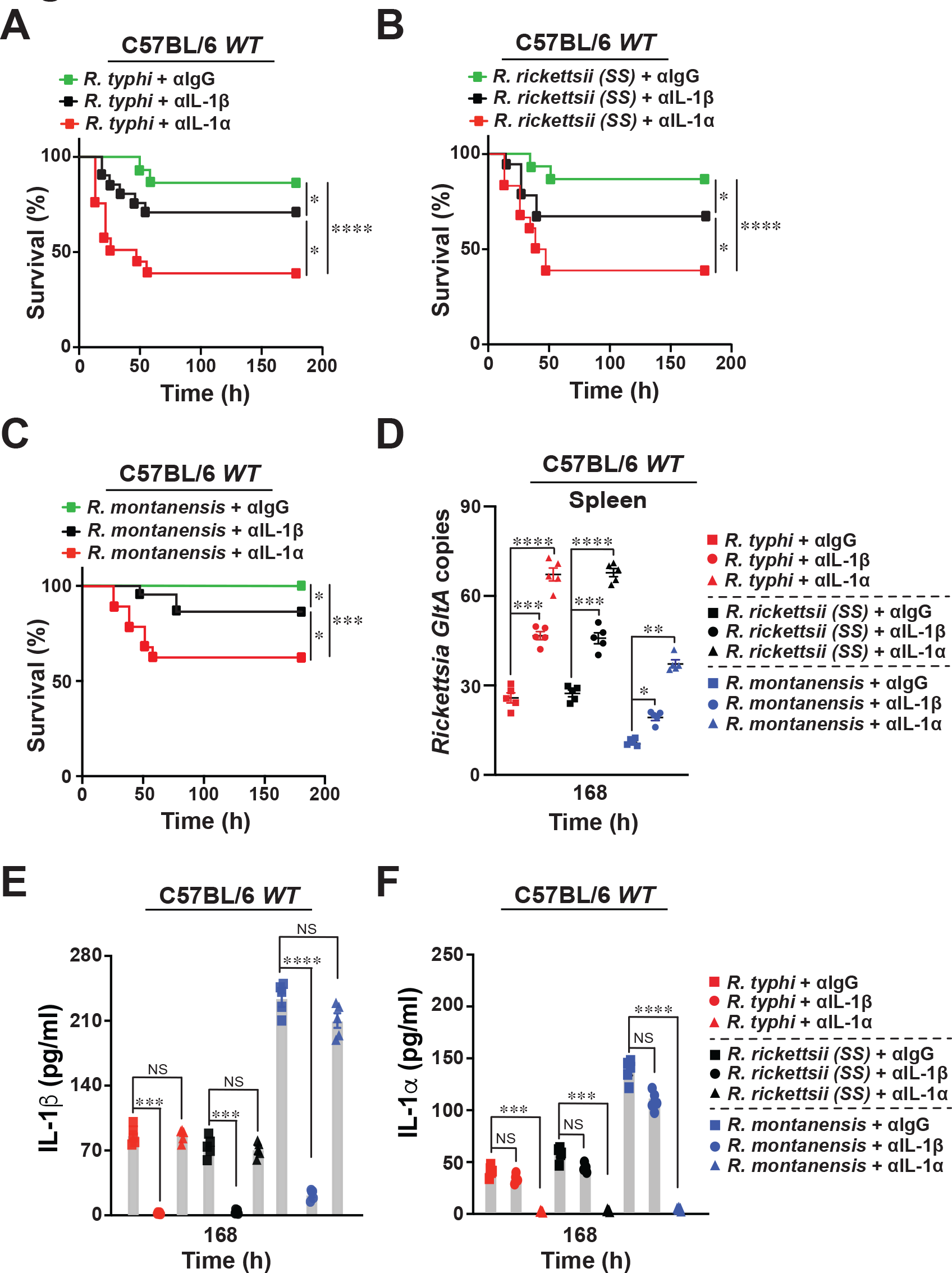
Neutralization of IL-1α activity augments mortality of pathogenic and non- pathogenic *Rickettsia*-induced rickettsiosis. (**A-C**) C57BL/6J *WT* mice were injected via tail vein (*i.v.*) with 10^5^ PFU of *R. typhi* [A], *R. rickettsii* [B], or *R. montanensis* [C] (A-C; n=12 for each treatment) followed by a subsequent injection (*i.v.*) with αIL-1β, αIL-1α or IgG-isotype control antibody (Ab) [250 μg Ab/mouse]. Survival was monitored for 7 days. (**D**) Bacterial burden was tested in spleens of Ab-treated *R. typhi*-, *R. rickettsii*-, and *R. montanensis*-injected *WT* mice shown in (A-C) by *GltA* RT-qPCR at day 7 (n=5 for each treatment) using the host housekeeping gene, *Gapdh*, for normalization. (**E-F**) Serum samples from mice described in (A-C) were analyzed for IL-1β (E) and IL- 1α (F) production at day 7 (n=5 for each treatment) using the Legendplex kits (BioLegend) followed by flow cytometry analysis. Error bars in (D-F) represent means ± SEM from five independent experiments; NS, non- significant; \**p* ≤ 0.05; \*\**p* ≤ 0.01; \*\*\**p* ≤ 0.005; \*\*\*\**p* ≤ 0.001.

**Fig. 3.**
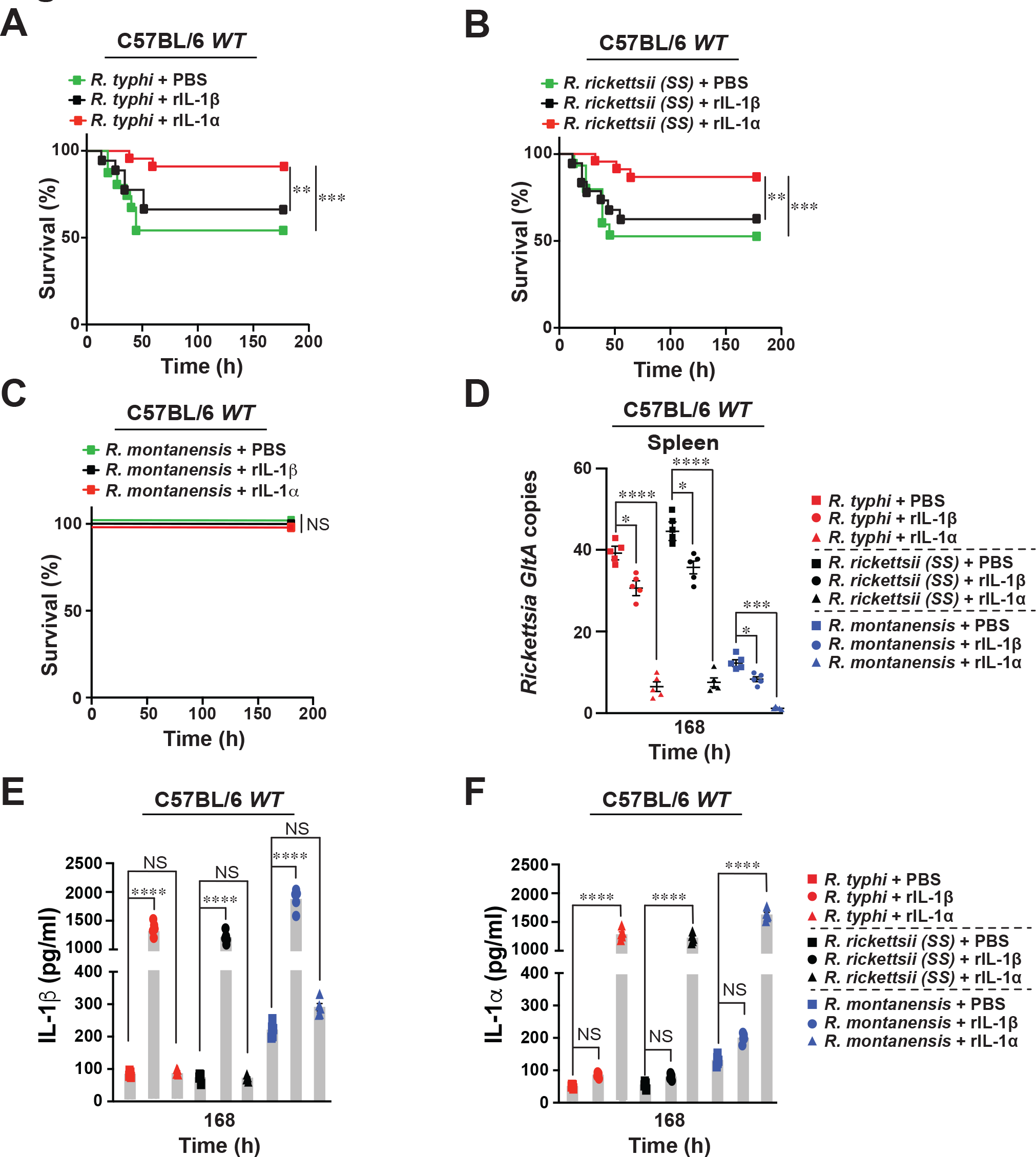
Administration of recombinant IL-1α rescues fatal *Rickettsia*-induced rickettsiosis. (**A-C**) C57BL/6J *WT* mice were injected *i.v.* with recombinant (r)IL-1β or rIL-1α protein [500 ng/mouse] followed by infection (24 h post protein injection) with 10^6^ PFU of *R. typhi* [A], *R. rickettsii* [B], *R. montanensis* [C], or PBS (A-C; n=15 for each treatment). Survival was monitored for 7 days. (**D**) Bacterial burden was tested in spleens of protein-treated *R. typhi*-, *R. rickettsii*-, *R. montanensis*- or PBS-injected *WT* mice shown in (A-C) by *GltA* RT-qPCR at day 7 (n=5 for each treatment) using the housekeeping gene, *Gapdh*, for normalization. (**E-F**) Serum samples from mice described in (A-C) were analyzed for IL-1β (E) and IL- 1α (F) production at day 7 (n=5 for each treatment) using the Legendplex kits (BioLegend) followed by flow cytometry analysis. Error bars in (D-F) represent means ± SEM from five independent experiments; NS, non- significant; \**p* ≤ 0.05; \*\**p* ≤ 0.01; \*\*\**p* ≤ 0.005; \*\*\*\**p* ≤ 0.001.

### Pathogenic, but not non-pathogenic, *Rickettsia* species block IL-1α secretion and avoid pro-IL-1α induction to establish a replication niche in macrophages

As MΦ are one of the cell types first encountered during infection by *Rickettsia* spp. and are considered to play a crucial role in either terminating the infection early at the skin site or allow initial pathogen colonization and subsequent dissemination within the infected host (3), we tested the hypothesis that pathogenic, but not non-pathogenic, rickettsiae suppress MΦ immune responses. In this effort, we determine the importance of IL-1α and IL-1β in restricting the replication of *Rickettsia* spp. by infecting BMDMΦ derived from *WT*, *Il-1β^-/-^* or *Il-1α^-/-^* mice with pathogenic *R. typhi* and *R. rickettsii* or non-pathogenic *R. montanensis* and assessed the cytokine levels of IL-1β and IL-1α in cultured supernatants as well as bacterial burdens. Infection of *Il-1β^-/-^* or *Il-1α^-/-^* BMDMΦ with *Rickettsia* spp. did not result in the secretion of either IL-1β or IL-1α, respectively, when compared to infected *WT* BMDMΦ (Figs. 4A-F). Moreover, infections with *R. montanensis* resulted in overall higher IL-1 responses and lower bacterial burden when compared to infections performed with *R. typhi* or *R. rickettsii* in *WT*, *Il-1β^-/-^* or *Il-1α^-/-^* BMDMΦ (Figs. 4A-G). Of note, the bacterial burden of all three *Rickettsia* spp. in infected *Il-1α^-/-^* BMDMΦ was higher than the levels detected in *Il-1β^-/-^* or *WT* BMDMΦ (Fig. 4G), suggesting that IL-1α and to significantly lesser extent IL-1β, plays a role in restricting rickettsiae survival. In agreement with our *in vivo* data, infection assays using BMDMΦ also displayed an overall higher bacterial burden of both pathogenic *Rickettsia* spp., as compared to the non-pathogenic *Rickettsia* strain (Figs. 4G and 1E). We further tested the protein expression levels of pro-IL-1β and -IL-1α upon bacterial-infection of *WT* BMDMΦ and showed that pro-IL-1β levels were induced by all three *Rickettsia* spp. to similar levels (∼5-fold) when compared to uninfected *WT* BMDMΦ (Fig. 4H). Intriguingly, only *R. montanensis*-infected *WT* BMDMΦ produced significantly higher levels of pro-IL- 1α, as compared to *R. typhi*- or *R. rickettsii*-infected MΦ (Fig. 4H).

**Fig. 4.**
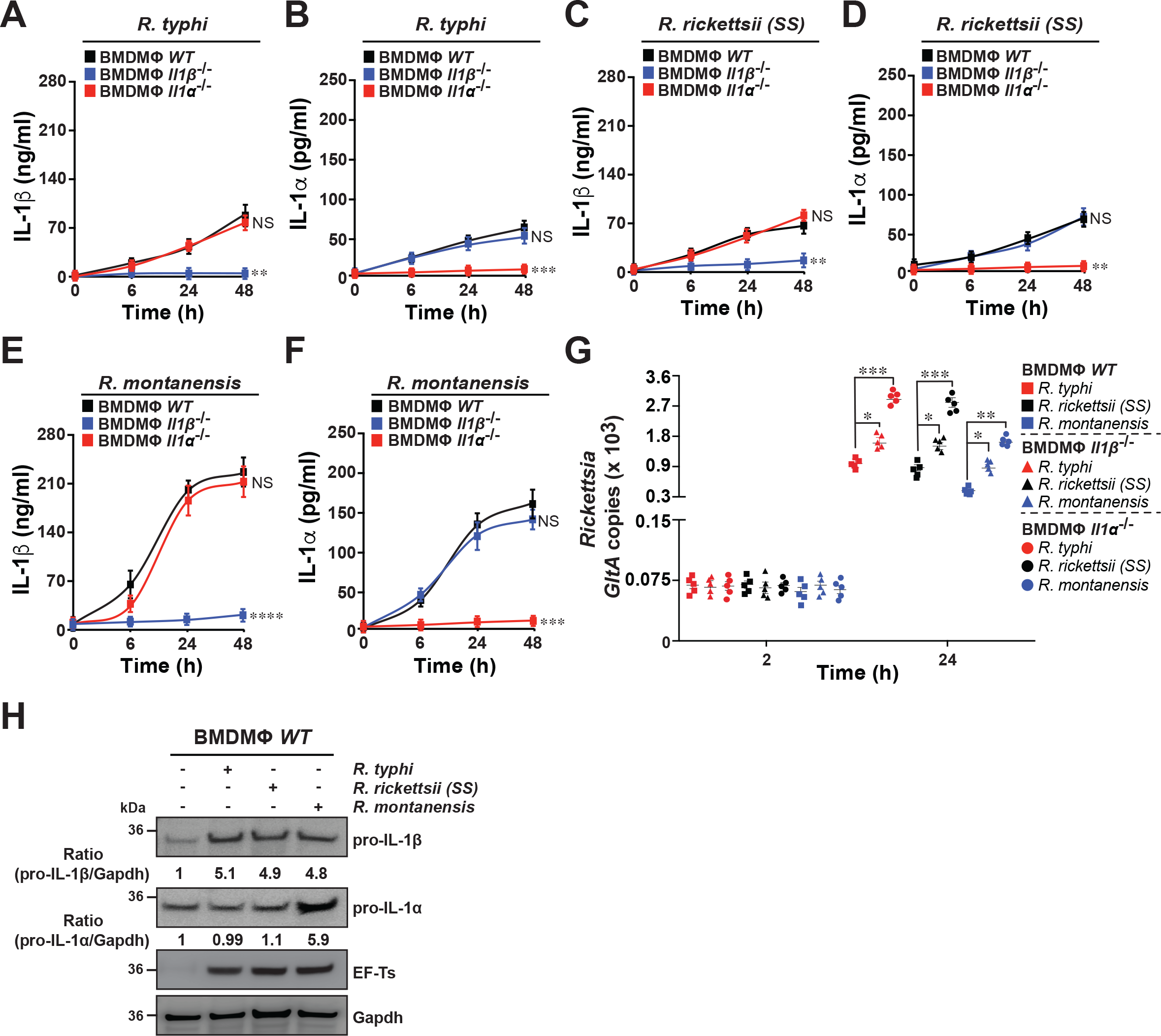
IL-1α, but not IL-1β, contributes to the survival of *Rickettsia* species in macrophages. (**A**-**F**) BMDMΦ from *WT*, *Il-1β^-/-^* or *Il-1α^-/-^* mice were infected with *R. typhi*, *R. rickettsii*, or *R. montanensis* [MOI=50] for 0, 6, 24, and 48 h. Culture supernatants were analyzed for production of IL-1β (A, C, and E) and IL-1α (B, D, and F) using Legendplex kits (BioLegend) followed by flow cytometry. (G) Bacterial burden in rickettsiae-infected BMDMΦ from *WT*, *Il-1β^-/-^* or *Il-1α^-/-^* mice was evaluated at 2 and 24 h post-infection by *GltA* RT-qPCR. Expression of the housekeeping gene *Gapdh* was used for normalization. (H) BMDMΦ from *WT* mice were either left uninfected (-) or infected with *R. typhi*, *R. rickettsii*, or *R. montanensis*, [MOI=50] for 24 h. Lysates were immunoblotted with αIL-1α, αIL-1β, αET-Ts, and αGapdh Abs. Densitometry was performed using Fiji software and data represent the fold-change ratios of pro-IL-1β/Gapdh or pro-IL-1α/Gapdh between uninfected and infected cells are shown. Immunoblot data are a representative of three independent experiments. Error bars in (A-G) represent means ± SEM from five independent experiments; NS: non- significant; \**p* ≤ 0.05; \*\**p* ≤ 0.01; \*\*\**p* ≤ 0.005; \*\*\*\**p* ≤ 0.001.

### Intracytosolic replication of pathogenic *Rickettsia* species in macrophages depends on the inhibition of IL-1 cytokine secretion via a Caspase-11-Gsdmd- dependent pathway

As our findings suggest that pathogenic, but not non-pathogenic, *Rickettsia* spp. prevent the activation of signaling pathways required for IL-1α and IL-1β production and release, we explored the mechanism of IL-1 signaling to a greater detail. As IL-1 signaling responses commonly involves the canonical and non-canonical inflammasome pathways, which in turn involves the proteolytic processing of both cytokines by activated Caspase (Casp)-1 (canonical) and Casp-11 (non-canonical) respectively, we assessed their potential role in regulating replication of non-pathogenic *vs*. pathogenic *Rickettsia* in BMDMΦ. In this effort, BMDMΦ derived from *WT*, *Casp-1^-/-^*, *Casp-11^-/-^*, and *Casp-1/11^-/-^* mice were infected with *R. typhi*, *R. rickettsii*, and *R. montanensis* and the levels of IL-1β and IL-1α, cell death, as well as bacterial burdens, were evaluated over the course of infection. Our assays revealed that Casp-11 was involved in the secretion of cleaved IL- 1α upon infection of BMDMΦ with non-pathogenic and pathogenic *Rickettsia* spp. (Figs. 5A-B, D-E, and G-H). In addition, Casp-11-deficiency resulted in a significant decrease in host cell death (Figs. 5C, F, and I). Casp-1-deficiency (*Casp-1^-/-^*) caused a significant decrease in IL-1β production [Figs. 5A, D, and G (green lines)], while IL-1α secretion as well as the level of cell death remained unaffected during infection with all three *Rickettsia* spp. [Figs. 5B-C, E-F, and H-I (green lines, symbols)]. Analysis of bacterial burdens further revealed a prominent role for Casp-11, but not for Casp-1, in restricting the replication of both pathogenic and non-pathogenic *Rickettsia* spp. (Fig. 5J; *WT* (square symbols); *Casp-1^-/-^* (sphere symbols); *Casp-11^-/-^* (triangles symbols); *Casp-1/11^-/-^* (diamond symbols)]. As our findings indicate that infections with *R. typhi*- and *R. rickettsii* resulted in a significant reduction of IL-1α secretion, likely via Casp-11-dependent mechanism, we assessed the expression and activation status of Casp-1 and Casp-11 via Western blot analyses. *R. montanensis* infection resulted in a robust activation of Casp-1, as indicative of the detection of the Casp-1-p20 fragment (Fig. 5K). In contrast, infection with *R. typhi* and *R. rickettsii* spp. resulted in a lower activation of Casp-1 (∼5- fold) (Fig. 5K). Intriguingly, only infection with *R. montanensis* resulted in a robust induction of Casp-11 (∼8 fold) as compared to infection data using both pathogenic *Rickettsia* spp. (Fig. 5K). To test if IL-1 cytokine secretion dependent on the bacterial load, we heat-inactivated both pathogenic and non-pathogenic *Rickettsia* spp. and showed that IL-1β and IL-1α release was significantly impaired as compared to infections using viable *Rickettsia* spp., a phenotype more strongly observed in infections using *R. montanensis* (Fig. S4). As IL-1 cytokine secretion dependents on Gsdmd, the pore-forming executor of pyroptosis (29–32), we assessed the proteolytic processing of Gsdmd and showed that only *R. montanensis* infection resulted in a robust cleavage of Gsdmd (∼8 fold), as indicative by the detection of the Gsdmd-p30 fragment (Fig. 5K). In support of our findings, we showed that infection of *Gsdmd^-/-^* BMDMΦ with non-pathogenic and pathogenic *Rickettsia* spp. released significantly lower levels of IL-1β and IL-1α, as compared to *WT* BMDMΦ (Figs. S5A-B). Furthermore, analysis of bacterial burdens provided additional evidence that Gsdmd plays a role in restricting the replication of *R. typhi*, *R. rickettsii*, and *R. montanensis* (Fig. S5C). These findings suggest that pathogenic, unlike non-pathogenic, *Rickettsia* spp. suppress IL-1 cytokine secretion via a Casp-11-Gsdmd-dependent pathway to support an intracytosolic replication in MΦ.

**Fig. 5.**
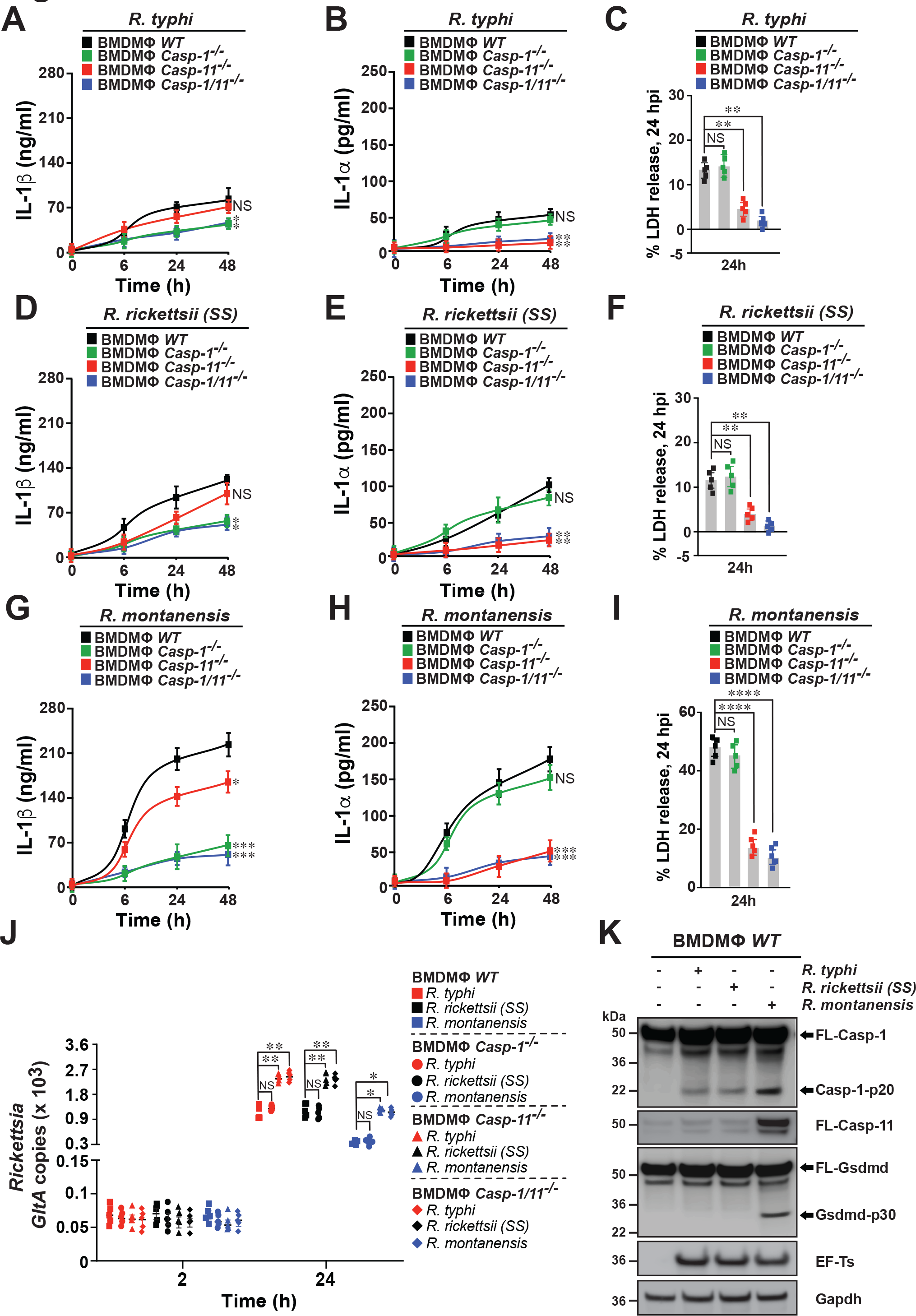
Pathogenic, but not non-pathogenic, *Rickettsia* species limit Caspase-1- and Caspase-11-dependent IL-1 signaling to facilitate their intracellular replication in macrophages. (**A-I**) BMDMΦ from *WT*, *Casp-1^-/-^*, *Casp-11^-/-^*, or *Casp-1/11^-/-^* mice were infected with *R. typhi* (A-C), *R. rickettsii* (D-F) or *R. montanensis* (G-I) [MOI=50] for 0, 6, 24, and 48 h. Culture supernatants were analyzed for production of IL-1β (A, D, G) and IL-1α (B, E, H) using Legendplex kits (BioLegend) followed by flow cytometry. BMDM cell death at 24 h post-infection, was measured by LDH release assay (C, F, I). (J) Bacterial burdens in infected BMDMΦ were evaluated 2 and 24 h post-infection by *GltA* RT-qPCR. Expression of the host housekeeping gene *Gapdh* was used for normalization. (K) Western analysis of Casp-1, Casp-11, and Gsdmd induction and processing at 24 h post-infection with *R. typhi*, *R. rickettsii*, or *R. montanensis* using αCasp-1, αCasp-11, and αGsdmd Abs. Reblotting with rickettsial specific αEF-Ts and host cell specific αGapdh Abs served as infection and equal loading controls respectively. Densitometry was performed using Fiji software and data represent the fold-change ratios of Casp-1-p20/ FL-Casp-1; Gsdmd-p30/FL-Gsdmd; or Casp-11/Gapdh between uninfected and infected cells are shown. Error bars in (A-J) represent means ± SEM from five independent experiments; NS: non- significant; \**p* ≤ 0.05; \*\**p* ≤ 0.01; \*\*\**p* ≤ 0.005; \*\*\*\**p* ≤ 0.001. Immunoblot data are a representative of three independent experiments.

### Secretion of IL-1α by macrophages is crucial in restricting the replication of pathogenic and non-pathogenic *Rickettsia* species *in vivo*

To examine whether secretion of IL-1 by MΦ limits the replication of *Rickettsia* spp. *in vivo*, we first injected (*i.v.*) *WT* mice with PBS- or dichloromethylene biphosphate (Cl2MBP)-liposomes to deplete endogenous macrophages as described previously [37]. Next, PBS- or Cl2MBP-liposomes treated *WT* mice were injected (*i.v.*) with BMDMΦ isolated from *WT*, *Il-1β^-/-^* or *Il-1α^-/-^* mice prior to infection with *R. typhi*, *R. rickettsii*, or *R. montanensis*. Strikingly, adoptive transfer of *Il-1α^-/-^* BMDMΦ, but not *Il-1β^-/-^*, or *WT* MΦ, significantly increased the mortality of Cl2MBP- but not PBS-treated *WT* mice injected with all three *Rickettsia* spp., reaching levels similar to the survival percentages observed in IL-1α Ab-neutralization studies (Figs. 6A-D, Fig. S6, and Fig. 2). Moreover, transfer of *Il-1α^-/-^* BMDMΦ resulted in the development of splenomegaly (Fig. S7) and an increase in bacterial burden in the spleens of Cl2MBP-treated *WT* mice (Fig. 6E), which correlated with a decrease in IL-1α serum concentrations without affecting IL-1β serum levels (Figs. 6F-G). In contrast, transfer of *Il-1β^-/-^* BMDMΦ, had an overall lesser effect on the severity of rickettsiosis, as evidenced by a lower mortality rate, smaller spleen size, and lower bacterial burden in Cl2MBP-treated *WT* mice (Figs. 6A-E, and Fig. S7), which is in agreement with our IL-1β Ab-neutralization data (Fig. 2). In addition, transfer of *Il-1β^-/-^* BMDMΦ, resulted in a decrease in IL-1β serum concentrations without affecting IL-1α serum levels (Figs. 6F-G). Collectively, these data suggest that modulation of expression and secretion of IL-1α secretion by macrophages is important to limit the replication of *Rickettsia* spp. *in vivo*.

**Fig. 6.**
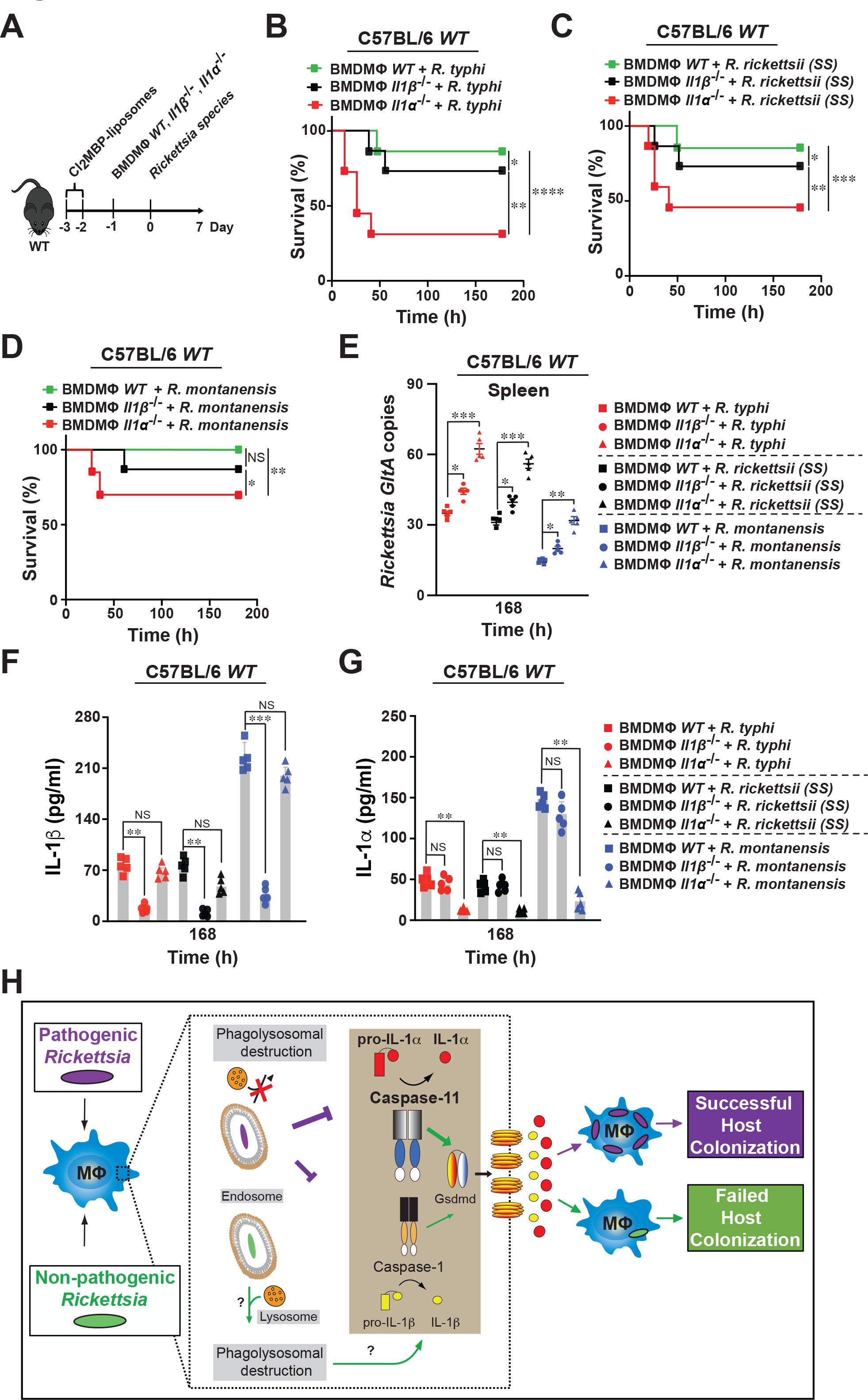
Macrophages-dependent secretion of IL-1α contributes more than IL-1β for controlling the survival and colonization of pathogenic and non-pathogenic *Rickettsia* species. (**A-D**) Dichloromethylene biphosphate (Cl2MBP)-treated C57BL/6J *WT* mice were injected *i.v.* with *WT*, *Il-1β^-/-^* or *Il-1α^-/-^* BMDMΦ [5 x 10^6^ cells/mouse] followed by infection (24 h post-MΦ transfer) with *R. typhi* [B], *R. rickettsii* [C], or *R. montanensis* [D] (dose 10^5^ PFU) (B-D; n=12 for each treatment). Survival was monitored for 7 days. (**E**) Bacterial burden was tested in spleens from mice described in (B-D) by *GltA* RT- qPCR at day 7 (n=5 for each treatment) using the housekeeping gene, *Gapdh*, for normalization. (**F-G**) Serum samples from mice described in (B-D) were analyzed for IL-1β (F) and IL- 1α (G) production at day 7 (n=5 for each treatment) using the Legendplex kits (BioLegend) followed by flow cytometry analysis. Error bars in (E-G) represent means ± SEM from five independent experiments; NS, non- significant; \**p* ≤ 0.05; \*\**p* ≤ 0.01; \*\*\**p* ≤ 0.005; \*\*\*\**p* ≤ 0.001. (**H**) Proposed working model on how pathogenic *Rickettsia* spp. suppress Casp-1 and Casp-11-dependent IL-1 signaling responses to establish a replication niche *in vitro* and *in vivo*. Of note, the majority of non-pathogenic *Rickettsia* spp. is likely destroyed by phagolysosomal fusion, while a subpopulation may escape lysosomal fusion, ultimately allowing for the induction of inflammasome mediated IL-1 responses.

## Discussion

Obligate intracellular bacterial pathogens, which successfully reside and replicate within the host cell, overcome responses of innate immune defense surveillance (*e.g.*, inflammasomes and autophagy). However, in the case of strict obligate intracytosolic *Rickettsia* spp., the roles for both inflammasome and autophagy to restrict their replication in endothelial cells and immune cells, like MΦ, is only now emerging, although without consistent mechanistical insights (23–28, 33). Given these knowledge gaps and our current lack of understanding on how TG *Rickettsia* spp. evade immune defense responses to facilitate host colonization, we established an animal model of rickettsiosis using C57BL/6J *WT* mice by comparing both mild (∼LD25) and more severe disease (∼LD50) development for two pathogenic spp., *R. rickettsii*, and *R. typhi*. In fact, data of disease severity correlated with the levels of bacterial burden detected in the mice spleens. We also observed that infections with pathogenic, but not non-pathogenic, *Rickettsia* spp. resulted in a reduced serum response of both pro-inflammatory cytokines, IL-1β and IL-1α, which is likely the result of pathogenic *Rickettsia* spp. to evade canonical and non-canonical inflammasome-dependent defense sensing, of which the former is in agreement with recent studies using *R. australis* (26). Thus, our data suggests that inhibition of both canonical and non-canonical inflammasome-dependent signaling contributes to the enhanced survival and colonization of pathogenic *Rickettsia* spp. in our *in vivo* experiments.

Our data further suggest that IL-1α, and to significantly lower extent IL-1β, plays a role in limiting rickettsial infection *in vivo*. By testing the putative anti-rickettsial capabilities of these cytokines through employing Ab-neutralization and recombinant protein assays, we showed that IL-1α, and to a much lesser level IL-1β, was able to restrict the replication and colonization of both non-pathogenic and pathogenic *Rickettsia* spp. *in vivo*. Our current understanding by which *Rickettsia* spp. evade host-induced anti-bacterial activities, and in particular, how cytosolic rickettsiae overcome host immune surveillance in defense cells, like MΦ, is primarily based on reports that are not aligned with one another (23–25, 27). Thus, we sought to address the hypothesis that pathogenic *R. typhi*, and *R. rickettsii* spp. but not non-pathogenic *R. montanensis* evade innate immune defense responses, in order, to establish an intracytosolic replication niche in MΦ. In agreement with our *in vivo* infection models, *R. montanensis*-infected *WT* BMDMΦ produced higher levels of IL-1α and IL-1β cytokines and displayed reduced bacterial loads during the course of infection, compared to *R. typhi*- or *R. rickettsii*-infected BMDMΦ. These data support the notion that non-pathogenic, but not pathogenic, *Rickettsia* spp. are more efficiently cleared by MΦ, a mechanism that further supports the previously published findings with SFG *Rickettsia* using THP-1 cells, a human macrophage-like cell line (34). Collectively, our presented data strengthen our hypothesis that pathogenic, but not non-pathogenic, *Rickettsia* spp. suppress anti-rickettsial inflammasome-dependent IL-1 cytokine responses to establish an intracytosolic replication niche in MΦ.

Given that IL-1 signaling is modulated through inflammasome-dependent Casp-1, Casp-11, and Gsdmd activation, we tested the role of both caspases as well as Gsdmd, and found that non-pathogenic, but not pathogenic, *Rickettsia* spp. ensued Casp-1 activation and Casp-11 induction, which ultimately resulted in the proteolytic processing of Gsdmd and release of IL-1α and IL-1β cytokines. Intriguingly, the lack of Casp-11 induction by pathogenic, but not non-pathogenic, *Rickettsia* spp. suggest that the membrane-bound LPS of *R. typhi*, and *R. rickettsii* spp. is likely less immunogenic than that of *R. montanensis*, which is further supported by our recent reports (35, 36). Our findings further suggest that pathogenic, as compared to non-pathogenic, *Rickettsia* spp., benefit from evasion of the Casp-11-Gsdmd-IL-1α signaling axis to establish a replication niche in MΦ as evidenced by the increased replication in MΦ from *Casp-11^-/-^*, *Casp-1/11^- /-^*, or *Gsdmd^-/-^* mice compared to *WT* and *Casp-1^-/-^* BMDMΦ. Finally, we sought to determine the role of IL-1 cytokine responses by MΦ in restricting the replication of *Rickettsia* spp. and showed that transfer of *Il-1α^-/-^* BMDMΦ, and to much lesser extent the administration of *Il-1β^-/-^* BMDMΦ, exacerbated the disease progression in *WT* mice injected with either pathogenic or non-pathogenic *Rickettsia* spp. It is worth noting that the observed differences in IL-1α release could be partially attributed to alternative mechanisms, including the retainment of IL-1α in the cytosol or nucleus, the dependance of IL-1β, and/or the association with the decoy receptor of IL-1 (IL-1R2), and future experiments are underway to address these possibilities (15, 29, 37–40). Also, IL-1α is produced by other immune cells, such as neutrophils (15). Although our study did not evaluate a potential contributing role of neutrophils, preceding findings suggest that neutrophils did not alter the course of rickettsiosis or contribute to the restriction of bacterial growth (28).

Importantly, preceding findings suggest that intracellular pathogens, like rickettsiae, not only encounter inflammasome-dependent defense mechanisms but are also confronted by another cytosolic defense pathway, autophagy (23, 25). Both responses not only are key to mount the appropriate host defense responses (16, 18) but are also functionally interconnected. In fact, recent reports indicated that autophagy acts on intracellular microbes upstream of the inflammasome and thereby functions as a negative regulator by degrading inflammasome components (10, 16, 18, 22). In the case of rickettsiae, however, the role of autophagy in regulating inflammasome responses to facilitate their host colonization remains inconclusive. For instance, *R. australis*, a pathogenic TRG member, benefited from ATG5-dependent autophagy induction and suppression of inflammasome-dependent IL-1β production to colonize MΦ (23, 25). In contrast, *R. parkeri*, a mildly pathogenic member of SFG, demonstrated that its surface protein OmpB is critical for protecting against autophagic recognition, while evasion of autophagy was critical for invasion of BMDMΦ and *WT* mice by *R. parkeri* (24, 27). Intriguingly, our recent report on *R. typhi* showed that *R. typhi* is ubiquitinated upon host entry, induces autophagy, but escapes autophagolysosomal maturation for intracellular colonization in non-phagocytic cells (9). Given these reports by others and our recent findings, it is tempting to speculate that pathogenic, but not non-pathogenic, *Rickettsia* spp. induce autophagy to down-regulate inflammasome-dependent IL-1β and IL-1α cytokine responses to establish an intracytosolic replication niche *in vitro* and *in vivo*, and our future research will address this possibility.

Overall, our findings present a previously unappreciated model of host invasion by which pathogenic, but not non-pathogenic, *Rickettsia* spp., avoid the activation of signaling pathways required for IL-1α production and release; likely via the suppression of the Casp-11-Gsdmd signaling pathway, to facilitate their intracytosolic replication in MΦ and ultimately cause host colonization (Fig. 6H).

## Materials and Methods

### Animals

All experiments were conducted in fully AAALAC accredited program using 8-10 weeks old female C57BL/6J *WT* mice in a specific pathogen-free environment according to the University of Maryland School of Medicine Institutional Animal Care and Use Committee (IACUC) in compliance with the National Institutes of Health Guide.

### Antibodies and reagents

Anti(α)-IL-1α (clone ALF-161), αIL-1β (clone B122), or an isotype control IgG (Armenian hamster IgG) antibody (Ab) were purchased from BioXCell. αCaspase (Casp)-1 Ab was purchased from Adipogen, while αCasp-11 [EPR18628] and αGsdmd [EPR19828] Abs were obtained from Abcam. Elongation factor Ts (EF-Ts) Ab was obtained from Primm Biotech as previously described (9), while the αGapdh (FL-335) Ab was purchased from Santa Cruz Biotechnology. Halt protease and phosphatase inhibitor cocktail was obtained from Thermo Fisher Scientific. Endotoxin-free recombinant mouse IL-1α and IL-1β proteins were purchased from BioLegend.

### Bacterial strains, cell culture, and infection

Vero76 cells (African green monkey kidney, ATCC, RL-1587) were maintained in minimal Dulbecco’s modified Eagle’s medium (DMEM) supplemented with 10% heat-inactivated fetal bovine serum (FBS) at 37°C with 5% CO2. *R. montanensis* strain M5/6 and *R. rickettsia* strain Sheila Smith (*SS*) were obtained from Dr. Ted Hackstadt (Rocky Mountain Laboratories, NIH, MT, USA) and *R. typhi* strain Wilmington was obtained from CDC. All *Rickettsia* strains were propagated in Vero76 cells grown in DMEM supplemented with 5% FBS at 34°C with 5% CO2. All *Rickettsia* were purified as previously described (7).

For infection of BMDMΦ, purified *Rickettsia* spp. were used at a multiplicity of infection (MOI) of 50, to ensure the presence of enough bacteria at early stage of infection, for host response (5, 8, 9). For infection using heat-inactivated bacteria, purified *Rickettsia* spp. were heated at 90°C for 20 min (41). Rickettsiosis in mice was induced by tail vein injection (*i.v.*) of purified *Rickettsia* (10^5^-10^6^ PFU), resuspended in PBS. At Days 1, 3, and 7 after administration, blood was collected, and serum cytokine levels were measured by flow cytometry. Splenic tissue specimens were collected at the indicated times and used for bacterial burden analysis by qPCR described below.

### Differentiation of bone marrow-derived macrophages

Bone marrow cells were isolated from femurs and tibias of *WT*, *Il-1β^-/-^* or *Il-1α^-/-^*, *Gsdmd^-/-^* , *Casp-1^-/-^*, *Casp-11^-/-^*, *Casp-1/Casp-11^-/-^* mice. Femurs from *Casp-1^-/-^*, *Casp-11^-/-^*, *Casp- 1/Casp-11^-/-^* were kindly provided by Dr. Amal Amer (The Ohio State University, OH, USA), while bones from *Il-1β^-/-^* or *Il-1α^-/-^* mice were obtained from Dr. Thirumala-Devi Kanneganti (St. Jude Children’s Research Hospital, TN, USA). Femurs from *Gsdmd^-/-^* were kindly provided by Dr. Matthew Welch (University of California, Berkeley, CA, USA). Differentiation was induced by culturing bone marrow cells in RPMI 1640 medium supplemented with 10% FBS and 30% L929-conditioned medium (a source of macrophage colony stimulating factor) and cultured for 7 days as described previously (42).

### Measurement of cytokines and chemokines

IL-1 cytokine concentrations in the sera of mice or supernatants from cultured BMDMΦ were assessed using the Legendplex mouse inflammation kit (BioLegend) following the manufacturer’s instructions as described previously (42).

### RNA isolation and quantitative real-time PCR

BMDMΦ samples were collected at 2, 6, 24, and 48 h post-infection, while spleens were collected at day 3 or 7 post infection. RNA was extracted from 1 x 10^6^ BMDMΦ or 100 μl of organ homogenate using the Quick-RNA miniprep kit (ZymoResearch). The iScript Reverse Transcription Supermix kit (Bio-Rad; 1708841) was used to synthesize cDNAs from 200 ng of RNA according to the manufacturer’s instructions. Quantitative real-time PCR was performed using SYBR Green (Thermo Fisher Scientific), 2 μl cDNA and 1 μM each of the following oligonucleotides for rickettsial citrate synthase gene (*gltA*) forward (F): 5’-CATAATAGCCATAGGATGAG-3’; reverse (R): 5’-ATGATTTATGGGGAACTACC-3’ and analyzed as described previously (25). Oligonucleotides for *Gapdh* were obtained from Qiagen.

### Extract preparation, and Western blot analysis

*Rickettsia*-infected BMDMΦ cells were lysed for 2 h at 4°C in ice-cold lysis buffer (50 mM HEPES [pH 7.4], 137 mM NaCl, 10% glycerol, 1 mM EDTA, 0.5% NP-40, and supplemented with protease and phosphatase inhibitory cocktails) as described previously (42). Equal amounts of protein were loaded for SDS-PAGE and membranes were probed with αCasp-1, αCasp-11, αGsdmd, αIL-1α, αIL-1β, αEF-Ts, and αGapdh Abs, followed by enhanced chemiluminescence with secondary Abs conjugated to horseradish peroxidase.

### Neutralization of endogenous IL-1α and IL-1β

For *in vivo* neutralization of IL-1α and IL-1β, C57BL/6J *WT* mice were *i.v.* injected with 250 mg of αIL-1α (clone ALF-161, BioXCell), αIL-1β (clone B122, BioXCell), or an IgG-isotype control (Armenian hamster IgG, BioXCell) Ab 24 h before the induction of mild rickettsiosis using 10^5^ PFU of *R. typhi*, *R. rickettsii*, or *R. montanensis*.

### Adoptive transfer of bone marrow-derived macrophages

C57BL/6J *WT* mice were injected (*i.v.*) twice with PBS- or dichloromethylene biphosphate (Cl2MBP)-liposomes 72 and 48 hours prior to macrophage transfer as described previously (42). Next, C57BL/6J *WT* mice were injected (*i.v.*) with *WT*, *Il-1β^-/-^* or *Il-1α^-/-^* BMDMΦ [5 x 10^6^ cells/mouse] followed by infection (24 hours post-MΦ transfer) with *R. typhi*, *R. rickettsii*, *R. montanensis*, or PBS (dose 10^5^ PFU).

### Statistical analysis

Endpoint studies of mice subjected to mild (10^5^ PFU) and severe (10^6^ PFU) rickettsiosis were analyzed by using Kaplan-Meir survival curves and the log-rank test (GraphPad Prism Software, version 8). The statistical significance was assessed using ANOVA with Tukey’s multiple comparisons post-test (GraphPad). Data are presented as mean ± SEM, unless stated otherwise. Alpha level was set to 0.05.

## Acknowledgments

We gratefully acknowledge Drs. Amal Amer (The Ohio State University, OH, USA), Thirumala-Devi Kanneganti (St. Jude Children’s Research Hospital, TN, USA), Matthew Welch (University of California, Berkeley, CA, USA) and Ted Hackstadt (Rocky Mountain Laboratories, NIH, MT, USA) for generously providing us with essential biological specimens and reagents, including femurs from various knock-out mice and rickettsial strains. We would like to thank Magda Beier-Sexton for her administrative, technical, organizational, and editorial contributions as well as Dr. Stefanie Vogel (University of Maryland School of Medicine, Baltimore, MD, USA) for critical discussions and editorial contributions to the manuscript. This work was supported with funds from the NIAID/NIH grants (R01AI017828 and R01AI126853 to A.F.A.).

---

**Fig. S1.**
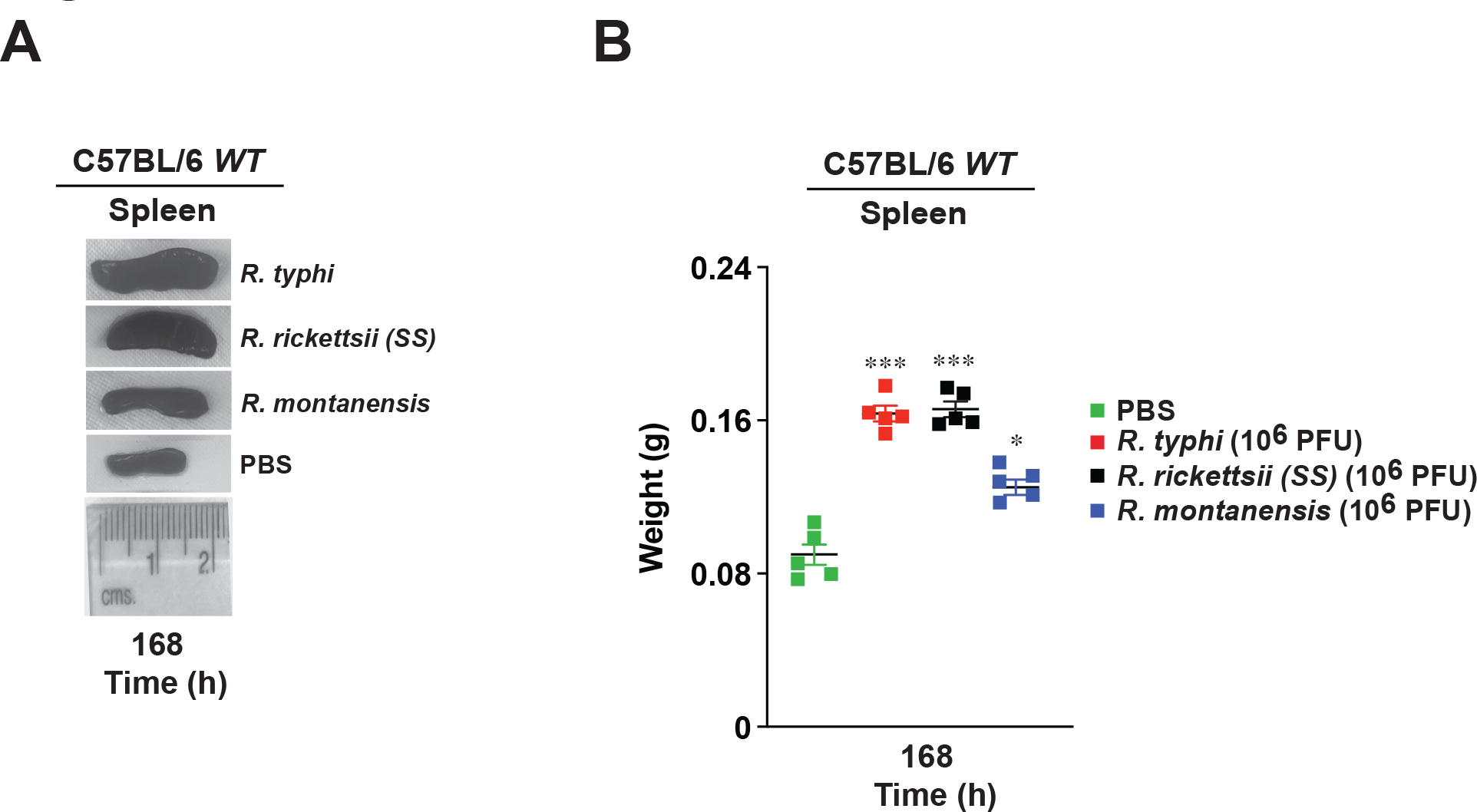
Splenic data during infection of pathogenic and non-pathogenic *Rickettsia* **species.** (A) Representative images of spleens (at day 7) from C57BL/6J *WT* mice injected *i.v*. with *R. typhi*, *R. rickettsii*, *R. montanensis*, or PBS (dose 10^6^ PFU). (B) Spleen weight from injected animals was evaluated at day 7 day (n=5). Error bars in (B) represent means ± SEM from five independent experiments; \**p* ≤ 0.05; \*\*\**p* ≤ 0.005.

**Fig. S2.**
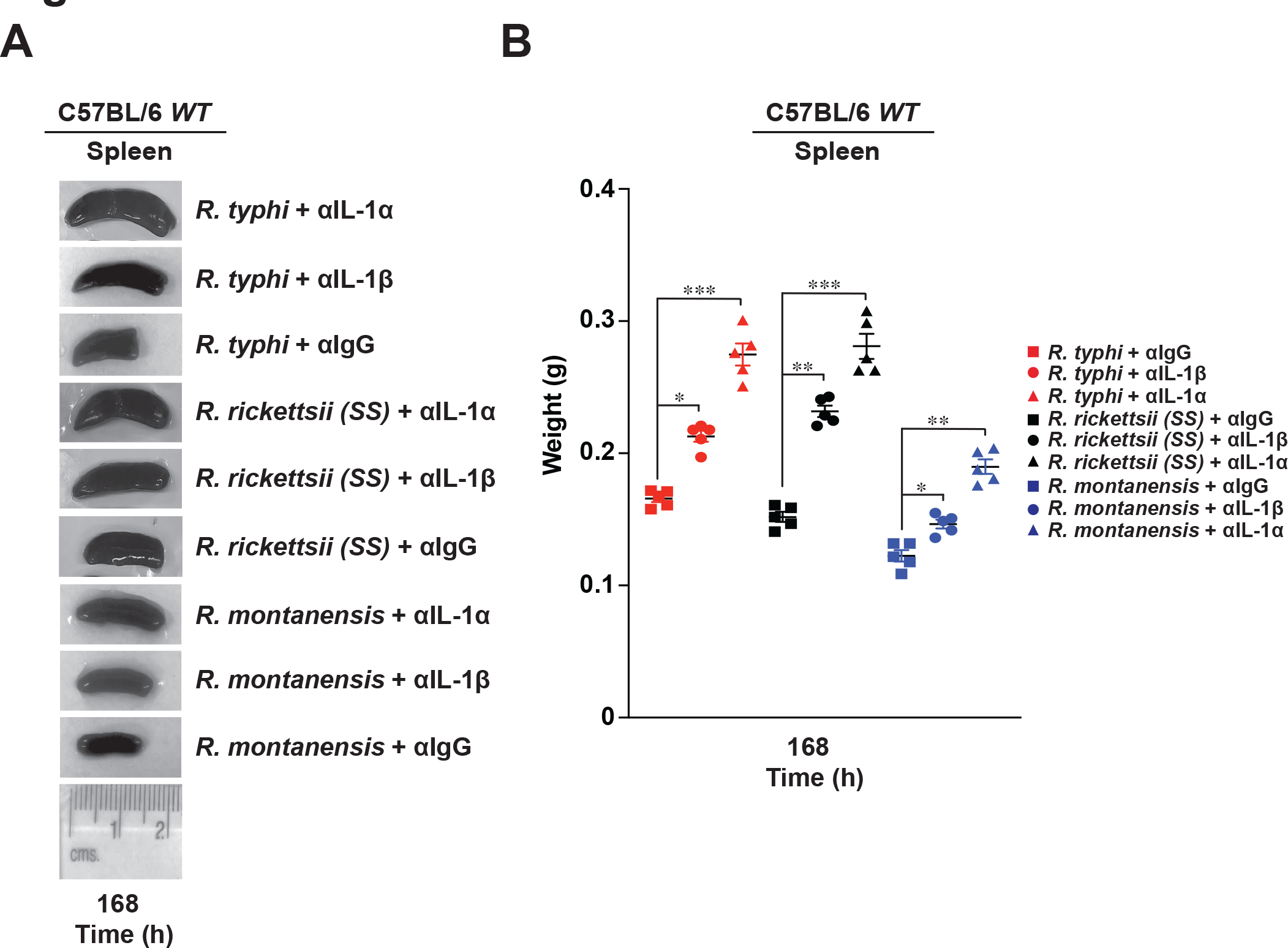
Splenic data after IL-1 signaling neutralization during infection of pathogenic and non-pathogenic *Rickettsia* species. (A) Representative images of spleens (day 7) from C57BL/6J *WT* mice injected *i.v.* with *R. typhi*, *R. rickettsii*, or *R. montanensis* (dose 10^5^ PFU) followed by administration of αIL- 1β, αIL-1α, or IgG isotype control antibody (Ab) [250 μg Ab/mouse] (24 h post-Ab injection). (B) Spleen weight from injected animals was evaluated at day 7 (n=5). Error bars in (B) represent means ± SEM from five independent experiments; \**p* ≤ 0.05; \*\**p* ≤ 0.01; \*\*\**p* ≤ 0.005.

**Fig. S3.**
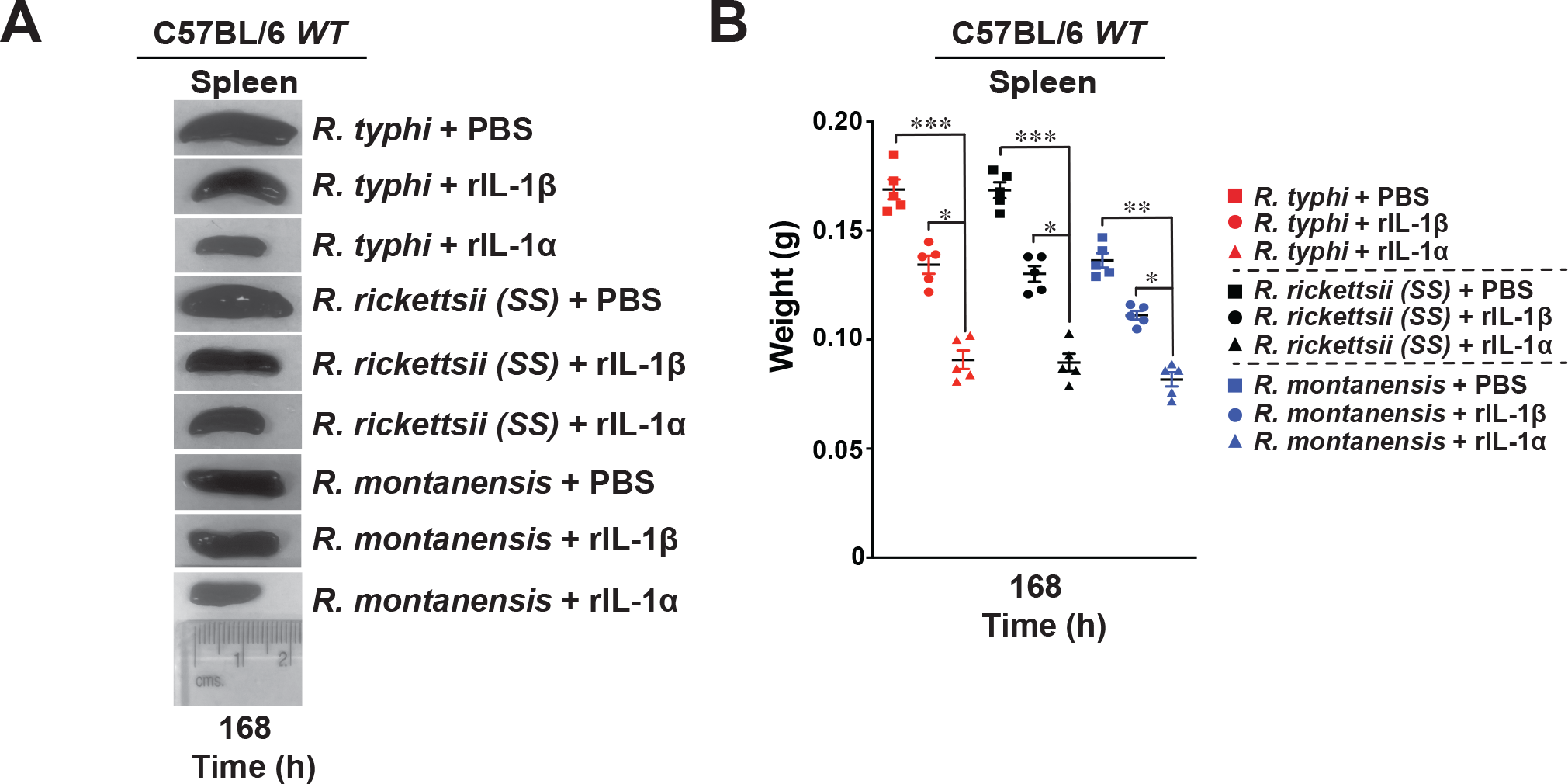
Splenic data after administration of recombinant IL-1α and IL-1β proteins during infection of pathogenic and non-pathogenic *Rickettsia* species. (A) Representative images of spleens (day 7) from C57BL/6J *WT* mice injected *i.v.* with rIL-1β or rIL-1α protein followed by administration of *R. typhi*, *R. rickettsii*, *R. montanensis* or PBS (dose 10^6^ PFU) at day 7. (B) Spleen weight from injected animals was evaluated at day 7 day (n=5). Error bars in (B) represent means ± SEM from five independent experiments; \**p* ≤ 0.05; \*\**p* ≤ 0.01; \*\*\**p* ≤ 0.005.

**Fig. S4.**
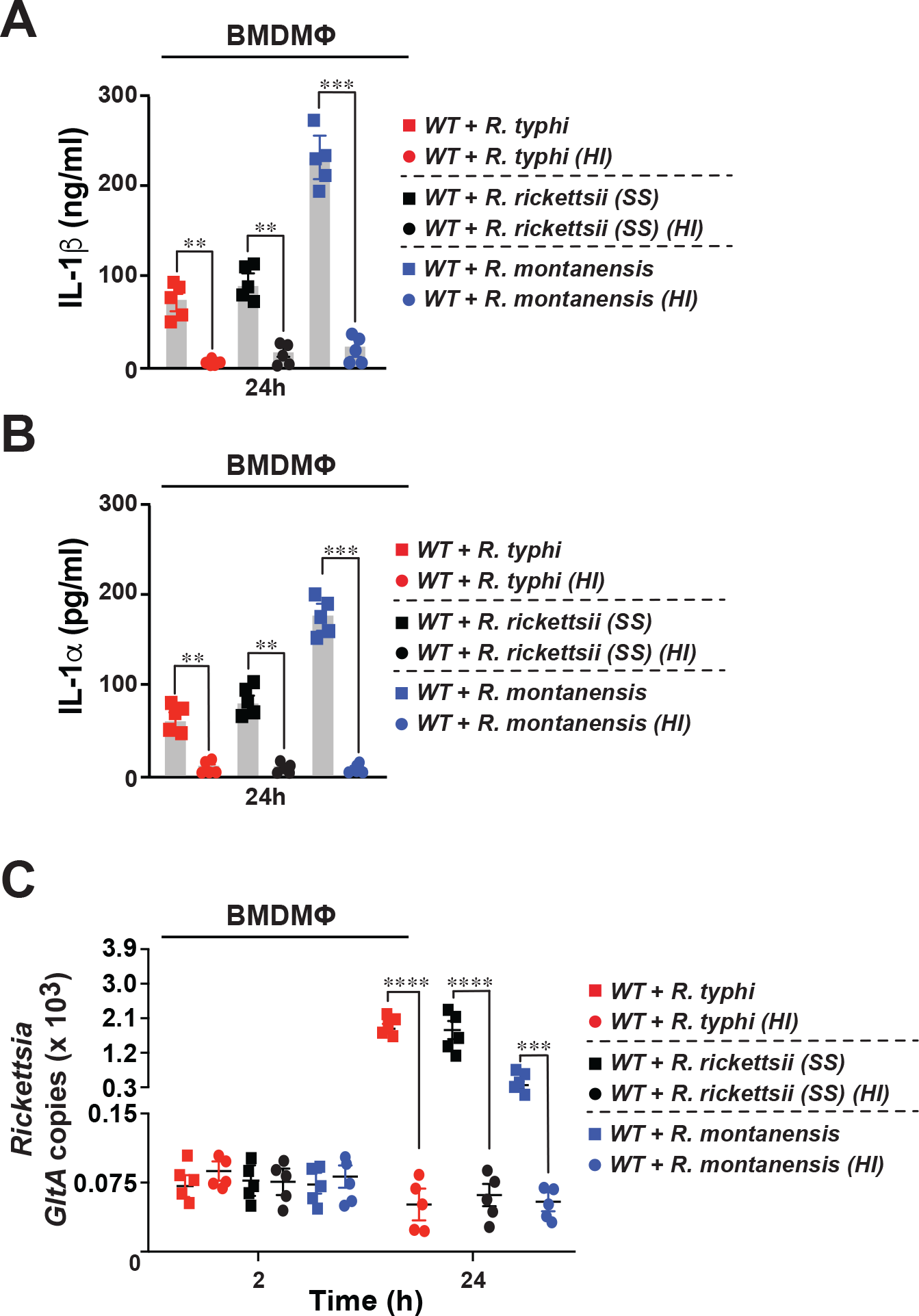
Heat-inactivation of *Rickettsia* species resulted in a reduced IL-1α and IL- 1β cytokine release by macrophages. (**A-B**) BMDMΦ from *WT* mice were infected with heat-inactivated or non-heat treated *R. typhi*, *R. rickettsii*, or *R. montanensis* [MOI=50] for 24 h. Culture supernatants were analyzed for production of IL-1β (A) and IL-1α (B) using Legendplex kits (BioLegend) followed by flow cytometry. (C) Bacterial burden in rickettsiae-infected BMDMΦ from *WT* mice was evaluated at 2 and 24 h post-infection by *GltA* RT-qPCR. Expression of the housekeeping gene *Gapdh* was used for normalization. Error bars in (A-C) represent means ± SEM from five independent experiments; NS: non- significant; \*\**p* ≤ 0.01; \*\*\**p* ≤ 0.005; \*\*\*\**p* ≤ 0.001.

**Fig. S5.**
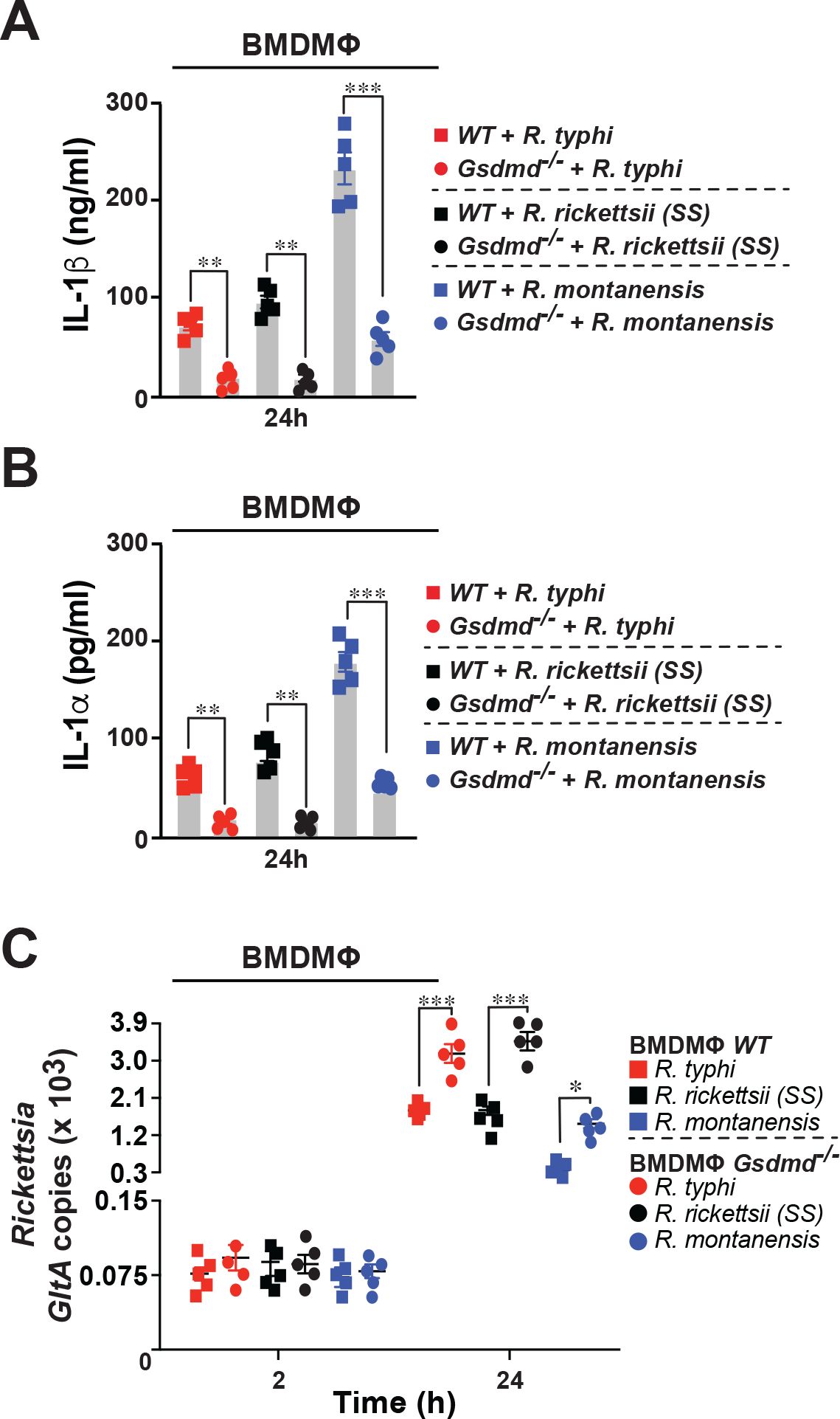
Gsdmd is involved in the release of IL-1α and IL-1β by macrophages upon ***Rickettsia* infection.** (**A-B**) BMDMΦ from *WT* and *Gsdmd^-/-^* mice were infected with *R. typhi*, *R. rickettsii*, or *R. montanensis* [MOI=50] for 24 h. Culture supernatants were analyzed for production of IL- 1β (A) and IL-1α (B) using Legendplex kits (BioLegend) followed by flow cytometry. (**C**) Bacterial burden in rickettsiae-infected BMDMΦ from *WT* and *Gsdmd^-/-^* mice was evaluated at 2 and 24 h post-infection by *GltA* RT-qPCR. Expression of the housekeeping gene *Gapdh* was used for normalization. Error bars in (A-C) represent means ± SEM from five independent experiments; NS: non- significant; \**p* ≤ 0.05; \*\**p* ≤ 0.01; \*\*\**p* ≤ 0.005.

**Fig. S6.**
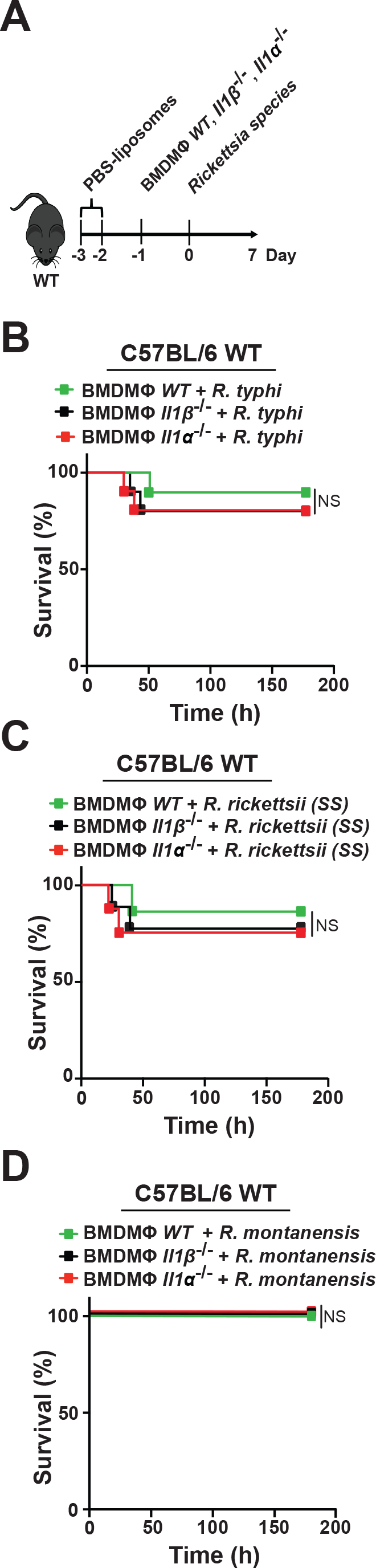
Survival data after adoptive transfer of *WT*, *Il-1β^-/-^* or *Il-1α^-/-^* macrophages and infection of pathogenic and non-pathogenic *Rickettsia* species into PBS- liposome-treated C57BL/6J *WT* mice. (A-D) PBS-treated C57BL/6J *WT* mice were injected *i.v.* with *WT*, *Il-1β^-/-^* or *Il-1α^-/-^* BMDMΦ [5 x 10^6^ cells/mouse] followed by infection (24 h post-MΦ transfer) with *R. typhi* [B], *R. rickettsii* [C], *R. montanensis* [D], or PBS (dose 10^5^ PFU) (B-D; n=12 for each treatment). Survival was monitored for 7 days.

**Fig. S7.**
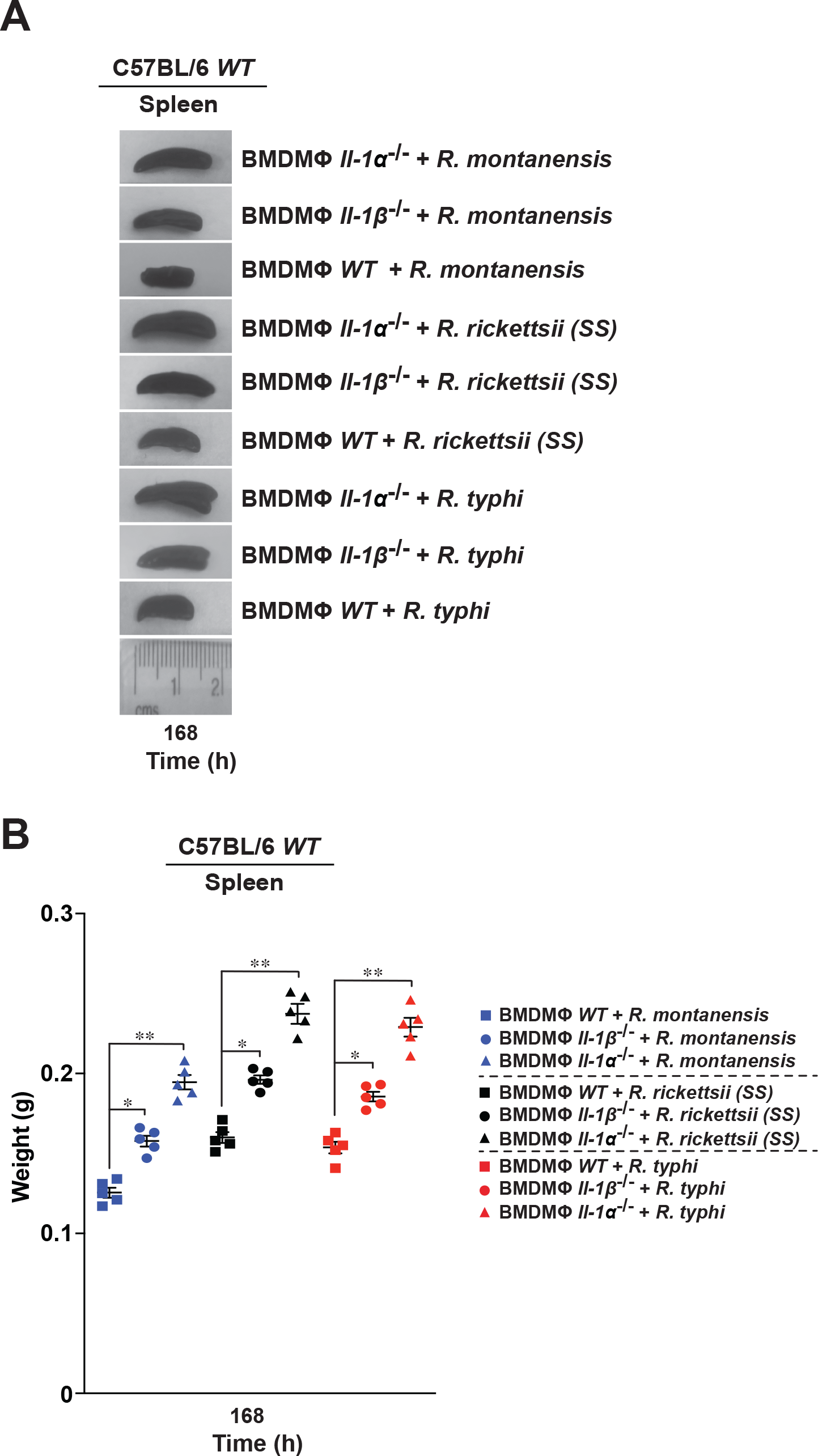
Splenic data after adoptive transfer of *WT*, *Il-1β^-/-^* or *Il-1α^-/-^* macrophages and infection of pathogenic and non-pathogenic *Rickettsia* species into dichloromethylene biphosphate (Cl2MBP)-treated C57BL/6J *WT* mice. (A) Representative images of spleens (day 7) from Cl2MBP-treated C57BL/6J *WT* mice injected *i.v.* with *WT*, *Il-1β^-/-^* or *Il-1α^-/-^* BMDMΦ [5 x 10^6^ cells/mouse] followed by administration (24 h post-MΦ transfer) of *R. typhi*, *R. rickettsii*, *R. montanensis*, or PBS (dose 10^5^ PFU). (B) Spleen weight from injected animals was evaluated at day 7 (n=5). Error bars in (B) represent means ± SEM; from five independent experiments; NS, non- significant; \**p* ≤ 0.05; \*\**p* ≤ 0.01.

## References

1. Ray K, Marteyn B, Sansonetti PJ, Tang CM. 2009. Life on the inside: the intracellular lifestyle of cytosolic bacteria. Nat Rev Microbiol 7:333–340.

2. Personnic N, Bärlocher K, Finsel I, Hilbi H. 2016. Subversion of Retrograde Trafficking by Translocated Pathogen Effectors. Trends Microbiol 24:450–462.

3. Sahni A, Fang R, Sahni SK, Walker DH. 2019. Pathogenesis of Rickettsial Diseases: Pathogenic and Immune Mechanisms of an Endotheliotropic Infection. Annu Rev Pathol 14:127–152.

4. Lehman SS, Noriea NF, Aistleitner K, Clark TR, Dooley CA, Nair V, Kaur SJ, Rahman MS, Gillespie JJ, Azad AF, Hackstadt T. 2018. The rickettsial ankyrin repeat protein 2 is a type IV secreted effector that associates with the endoplasmic reticulum. MBio 9.

5. Rahman MS, Ammerman NC, Sears KT, Ceraul SM, Azad AF. 2010. Functional characterization of a phospholipase A(2) homolog from Rickettsia typhi. J Bacteriol 192:3294–3303.

6. Rahman MS, Gillespie JJ, Kaur SJ, Sears KT, Ceraul SM, Beier-Sexton M, Azad AF. 2013. Rickettsia typhi Possesses Phospholipase A2 Enzymes that Are Involved in Infection of Host Cells. PLoS Pathog2013/07/03. 9:e1003399.

7. Rennoll-Bankert KE, Rahman MS, Gillespie JJ, Guillotte ML, Kaur SJ, Lehman SS, Beier-Sexton M, Azad AF. 2015. Which Way In? The RalF Arf-GEF Orchestrates Rickettsia Host Cell Invasion. PLoS Pathog 11:e1005115.

8. Rennoll-Bankert KE, Rahman MS, Guillotte ML, Lehman SS, Beier-Sexton M, Gillespie JJ, Azad AF. 2016. RalF-Mediated Activation of Arf6 Controls Rickettsia typhi Invasion by Co-Opting Phosphoinositol Metabolism. Infect Immun 84:3496– 3506.

9. Voss OH, Gillespie JJ, Lehman SS, Rennoll SA, Beier-Sexton M, Rahman MS, Azad AF. 2020. Risk1, a Phosphatidylinositol 3-Kinase Effector, Promotes Rickettsia typhi Intracellular Survival. MBio 11.

10. Sun Q, Fan J, Billiar TR, Scott MJ. 2017. Inflammasome and Autophagy Regulation: A Two-way Street. Mol Med 23:188–195.

11. Place DE, Kanneganti TD. 2018. Recent advances in inflammasome biology. Curr Opin Immunol. Elsevier Ltd.

12. Xue Y, Enosi Tuipulotu D, Tan WH, Kay C, Man SM. 2019. Emerging Activators and Regulators of Inflammasomes and Pyroptosis. Trends Immunol. Elsevier Ltd.

13. Wiggins KA, Parry AJ, Cassidy LD, Humphry M, Webster SJ, Goodall JC, Narita M, Clarke MCH. 2019. IL-1α cleavage by inflammatory caspases of the noncanonical inflammasome controls the senescence-associated secretory phenotype. Aging Cell 18:1–13.

14. Tapia VS, Daniels MJD, Palazón-Riquelme P, Dewhurst M, Luheshi NM, Rivers- Auty J, Green J, Redondo-Castro E, Kaldis P, Lopez-Castejon G, Brough D. 2019. The three cytokines IL-1β, IL-18, and IL-1α share related but distinct secretory routes. J Biol Chem 294:8325–8335.

15. Malik A, Kanneganti T-D. 2018. Function and regulation of IL-1α in inflammatory diseases and cancer. Immunol Rev 281:124–137.

16. Krakauer T. 2019. Inflammasomes, Autophagy, and Cell Death: The Trinity of Innate Host Defense against Intracellular Bacteria. Mediators Inflamm 2019:1–10.

17. Mitchell G, Isberg RR. 2017. Innate Immunity to Intracellular Pathogens: Balancing Microbial Elimination and Inflammation. Cell Host Microbe.

18. Seveau S, Turner J, Gavrilin MA, Torrelles JB, Hall-Stoodley L, Yount JS, Amer AO. 2018. Checks and Balances between Autophagy and Inflammasomes during Infection. J Mol Biol 430:174–192.

19. Storek KM, Monack DM. 2015. Bacterial recognition pathways that lead to inflammasome activation. Immunol Rev.

20. Vladimer GI, Marty-Roix R, Ghosh S, Weng D, Lien E. 2013. Inflammasomes and host defenses against bacterial infections. Curr Opin Microbiol.

21. Zheng D, Liwinski T, Elinav E. 2020. Inflammasome activation and regulation: toward a better understanding of complex mechanisms. Cell Discov 6:36.

22. Takahama M, Akira S, Saitoh T. 2018. Autophagy limits activation of the inflammasomes. Immunol Rev. Blackwell Publishing Ltd.

23. Bechelli J, Vergara L, Smalley C, Buzhdygan TP, Bender S, Zhang W, Liu Y, Popov VL, Wang J, Garg N, Hwang S, Walker DH, Fang R. 2019. Atg5 supports Rickettsia australis infection in macrophages in vitro and in vivo. Infect Immun 87.

24. Burke TP, Engström P, Chavez RA, Fonbuena JA, Vance RE, Welch MD. 2020. Inflammasome-mediated antagonism of type I interferon enhances Rickettsia pathogenesis. Nat Microbiol 5:688–696.

25. Smalley C, Bechelli J, Rockx-Brouwer D, Saito T, Azar SR, Ismail N, Walker DH, Fang R. 2016. Rickettsia australis Activates Inflammasome in Human and Murine Macrophages. PLoS One 11:e0157231.

26. Rumfield C, Hyseni I, McBride JW, Walker DH, Fang R. 2020. Activation of ASC inflammasome driven by toll-like receptor 4 contributes to host immunity against rickettsial infection. Infect Immun 88.

27. Engström P, Burke TP, Mitchell G, Ingabire N, Mark KG, Golovkine G, Iavarone AT, Rape M, Cox JS, Welch MD. 2019. Evasion of autophagy mediated by Rickettsia surface protein OmpB is critical for virulence. Nat Microbiol 4:2538– 2551.

28. Papp S, Moderzynski K, Rauch J, Heine L, Kuehl S, Richardt U, Mueller H, Fleischer B, Osterloh A. 2016. Liver Necrosis and Lethal Systemic Inflammation in a Murine Model of Rickettsia typhi Infection: Role of Neutrophils, Macrophages and NK Cells. PLoS Negl Trop Dis 10:e0004935.

29. Batista SJ, Still KM, Johanson D, Thompson JA, O’Brien CA, Lukens JR, Harris TH. 2020. Gasdermin-D-dependent IL-1α release from microglia promotes protective immunity during chronic Toxoplasma gondii infection. Nat Commun 11:1–13.

30. Kanneganti A, Malireddi RKS, Saavedra PHV, Walle L Vande, Van Gorp H, Kambara H, Tillman H, Vogel P, Luo HR, Xavier RJ, Chi H, Lamkanfi M. 2018. GSD MD is critical for autoinflammatory pathology in a mouse model of Familial Mediterranean Fever. J Exp Med 215:1519–1529.

31. Liu X, Zhang Z, Ruan J, Pan Y, Magupalli VG, Wu H, Lieberman J. 2016. Inflammasome-activated gasdermin D causes pyroptosis by forming membrane pores. Nature 535:153–158.

32. Tsuchiya K, Hosojima S, Hara H, Kushiyama H, Mahib MR, Kinoshita T, Suda T. 2021. Gasdermin D mediates the maturation and release of IL-1α downstream of inflammasomes. Cell Rep 34:108887.

33. Moderzynski K, Papp S, Rauch J, Heine L, Kuehl S, Richardt U, Fleischer B, Osterloh A. 2016. CD4+T Cells Are as Protective as CD8+T Cells against Rickettsia typhi Infection by Activating Macrophage Bactericidal Activity. PLoS Negl Trop Dis 10.

34. Curto P, Simões I, Riley SP, Martinez JJ. 2016. Differences in intracellular fate of two spotted fever group Rickettsia in macrophage-like cells. Front Cell Infect Microbiol 6.

35. Guillotte ML, Gillespie JJ, Chandler CE, Rahman MS, Ernst RK, Azad AF. 2018. Rickettsia lipid A biosynthesis utilizes the late acyltransferase LpxJ for secondary fatty acid addition. J Bacteriol 200.

36. Guillotte ML, Chandler CE, Verhoeve VI, Gillespie JJ, Driscoll TP, Rahman MS, Ernst RK, Azad AF. 2021. Lipid A Structural Divergence in Rickettsia Pathogens. mSphere 6.

37. Fettelschoss A, Kistowska M, LeibundGut-Landmann S, Beer H-D, Johansen P, Senti G, Contassot E, Bachmann MF, French LE, Oxenius A, Kündig TM. 2011. Inflammasome activation and IL-1β target IL-1α for secretion as opposed to surface expression. Proc Natl Acad Sci U S A 108:18055.

38. Paolo NC Di, Shayakhmetov DM. 2016. Interleukin 1α and the inflammatory process. Nat Immunol 17:906.

39. Peters VA, Joesting JJ, Freund GG. 2013. IL-1 receptor 2 (IL-1R2) and its role in immune regulation. Brain Behav Immun 32:1.

40. Shi L, Song L, Maurer K, Dou Y, Patel VR, Su C, Leonard ME, Lu S, Hodge KM, Torres A, Chesi A, Grant SFA, Wells AD, Zhang Z, Petri MA, Sullivan KE. 2020. IL-1 Transcriptional Responses to Lipopolysaccharides Are Regulated by a Complex of RNA Binding Proteins. J Immunol 204:1334–1344.

41. Ammerman NC, Beier-Sexton M, Azad AF. 2008. Laboratory Maintenance of Rickettsia rickettsii. Curr Protoc Microbiol Chapter 3:Unit 3A.5.

42. Voss OH, Murakami Y, Pena MY, Lee H-N, Tian L, Margulies DH, Street JM, Yuen PST, Qi C-F, Krzewski K, Coligan JE. 2016. Lipopolysaccharide-Induced CD300b Receptor Binding to Toll-like Receptor 4 Alters Signaling to Drive Cytokine Responses that Enhance Septic Shock. Immunity 44:1365–1378.

43. Voss OH, Arango D, Tossey JC, Villalona Calero MA, Doseff AI. 2021. Splicing reprogramming of TRAIL/DISC-components sensitizes lung cancer cells to TRAIL-mediated apoptosis. Cell Death Dis 12.

